# Discovery and Engineering of the l-Threonine Aldolase from *Neptunomonas Marine* for Efficient Synthesis of β-Hydroxy-α-Amino Acids via C–C Formation

**DOI:** 10.1101/2023.04.09.536162

**Authors:** Yuanzhi He, Siyuan Li, Jun Wang, Xinrui Yang, Jiawei Zhu, Qi Zhang, Li Cui, Zaigao Tan, Wupeng Yan, Yong Zhang, Luyao Tang, Lin-Tai Da, Yan Feng

## Abstract

l-Threonine aldolases (LTAs) are attractive biocatalysts for synthesizing β-hydroxy-α-amino acids (HAAs) via C–C bond formation in pharmaceuticals, although their industrial applications suffer from low activity and diastereoselectivity. Herein, we describe the discovery of a new LTA from *Neptunomonas marine* (*Nm*LTA) that displays both ideal enzymatic activity (64.8 U/mg) and diastereoselectivity (89.5% diastereomeric excess; de) for the desired product l-*threo*-4-methylsulfonylphenylserine (l-*threo*-MPTS). Using X-ray crystallography, site-directed mutagenesis, and computational modeling, we propose a “dual-conformation” mechanism for the diastereoselectivity control of *Nm*LTA, whereby the incoming 4-methylsulfonylbenzaldehyde (4-MTB) could potentially bind at the *Nm*LTA active site in two distinct orientations, potentially forming two diastereoisomers (*threo*- or *erythro*-form products). Importantly, two key *Nm*LTA residues H140 and Y319 play critical roles in fine-tuning the binding mode of 4-MTB, supported by our site-mutagenesis assays. Uncovering of the catalytic mechanism in *Nm*LTA guides us to further improve the diastereoselectivity of this enzyme. A triple variant of *Nm*LTA (N18S/Q39R/Y319L; SRL) exhibited both improved diastereoselectivity (de value > 99%) and enzymatic activity (95.7 U/mg) for the synthesis of l-*threo*-MPTS compared with that of wild type. The preparative gram-scale synthesis for l-*threo*-MPTS with the SRL variant produced a space-time yield of up to 9.0 g L^−1^h^−1^, suggesting a potential role as a robust C–C bond synthetic tool for industrial synthesis of HAAs at a preparative scale. Finally, the SRL variant accepted a wider range of aromatic aldehyde derivatives as substrates and exhibited improved diastereoselectivity toward *para*-site substituents. This work provides deep structural insights into the molecular mechanism underlying the catalysis in *Nm*LTA and pinpoints the key structural motifs responsible for regulating the diastereoselectivity control, thereby guiding future attempts for protein engineering of various LTAs from different sources.

## INTRODUCTION

Carbon–carbon (C–C) bond forming reactions are highly effective strategies for constructing complex molecules by connecting small building blocks.^1, 2^ The aldol reaction is an attractive approach to stereoselectively synthesize functionalized chiral amino acids via C–C formation.^3–6^ Enzymatic catalysis has been widely applied in aldol reaction for environmental-friendly and efficient synthesis of numerous valuable chemicals and pharmaceuticals by enabling ideal stereoselectivity.^7–9^ In particular, microbial l-threonine aldolases (LTAs), which are pyridoxal-5ʹphosphate (PLP)-dependent enzymes^10^, have received considerable attention for functionalizing the C–C bond by catalyzing the aldol reaction of various aldehydes and α-amino acids in one-step to form diverse β-hydroxy-α-amino acids (HAAs) with two adjacent chiral centers (C_α_ and C_β_) (Scheme 1).^1, 11–13^ HAAs belong to an important subclass of chiral intermediates with antimicrobial, antitumor, and immunosuppressive activities, and are therefore widely used to produce valuable pharmaceuticals (Figure S1).^14–18^ For example, l-*threo*-4-methylsulfonylphenylserine (l-*threo*-MPTS) serves as an important intermediate in the synthesis of the glycopeptide antibiotics florfenicol and thiamphenicol,^19–21^ and l-*threo*-4-nitrophenylserine is a key precursor for the production of chloramphenicol.^22^ Although LTA has strict stereoselectivity at the C_α_-site, the stereoselectivity at the C_β_-site is much less conserved.^23, 24^ However, initial efforts to improve the diastereoselectivity of *Pseudomonas* sp. LTA (*Ps*LTA) using error-prone PCR only had a limited effect on increasing the diastereomeric excess (de) value.^14, 25^

**Scheme 1.**
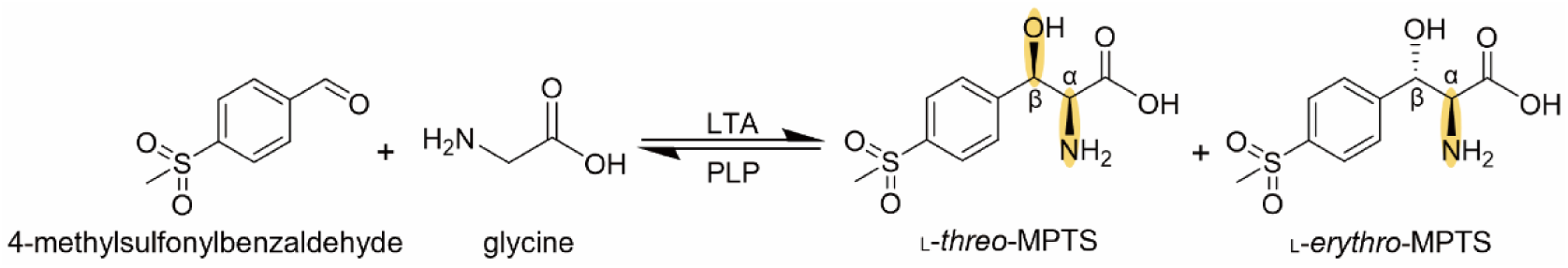
The aldol reaction catalyzed by LTA for the synthesis of _L_-*threo*-MPTS.

In comparison, structure-based engineering strategies have become more efficient owing to the availability of an increasing number of LTA structures from various sources.^26–29^ These structures reveal that LTA forms a tetrameric protein whose active-site residues are located within the interface between adjacent monomers.^28^ The LTA conformation was captured in various states, including the apo state, and in complexes with varied substrates in binary and ternary forms.^26–29^ Thus, these crystal structures provide atomic-level detail of the molecular recognitions between LTA and its substrates, thereby guiding further enzyme engineering. Several groups have designed a number of LTA variants with improved stereoselectivity at the C_β_-site based on these crystal structures, or homolog models.^14, 17, 30–32^ For example, Linʹs group increased the de value of *Actinocorallia herbida* LTA (*Ah*LTA) from 58% to 81% for l-*threo*-MPTS by modulating the substrate binding pocket,^19^ and they further enhanced the *Ah*LTA de value to 93.7% via rational design by combining molecular dynamics simulations and free-energy calculations.^19, 23^ Cai et al. improved the diastereoselectivity of *Pseudomonas putida* LTA (*Pp*LTA) from 32% to 85% for l-*threo*-4-nitrophenylserine via computer-aided directed evolution.^22^ In addition, Zheng and colleagues conducted a series of structure-based designs of LTA variants to produce an enhanced de value up to 99% for l-*threo*-MPTS.^31, 33^ This group also proposed that the substrate 4-methylsulfonylbenzaldehyde (4-MTB) could potentially bind to LTA along two distinct channels, enabling in the formation of two diastereomeric products.^30–33^

Although the diastereoselectivities of certain LTAs have been improved via protein engineering, their enzymatic activities were considerably undermined,^30–32^ which limited their applications in HAA-related pharmaceutical synthesis. Currently, it remains challenging to simultaneously improve the activity and C_β_-site stereoselectivity of LTAs. Moreover, given that the structure of the aldehyde-bound LTA complex is still unknown, the key structural motifs involved in regulating the LTA diastereoselectivity remain obscure, hampering further protein engineering. Intriguingly, the enzymes from marine microorganisms can exhibit remarkable activities owing to their habitat-related characteristics, such as using unique carbon sources, including petroleum, polycyclic aromatic hydrocarbons, and fatty acid residues from marine animals.^34–37^ Therefore, we sought to explore new LTAs from marine sources to mine highly active LTAs for C–C condensation with high diastereoselectivity as well as ideal activity.

Herein, we describe the discovery of a highly active *Neptunomonas marine* LTA (*Nm*LTA) that synthesized l-*threo*-MPTS with the highest enzymatic activity and diastereoselectivity among the currently reported native LTAs. By combining X-ray crystallography and computational modeling, we pinpointed the key regulatory motifs in *Nm*LTA responsible for controlling the diastereoselectivity of the desired product, which prompted us to propose a novel diastereoselectivity mechanism. Furthermore, site-directed and combinatorial mutagenesis assays identified a triple mutant of *Nm*LTA, N18S/Q39R/Y319L (termed SRL), that could further improve the de value > 99% for l-*threo*-MPTS, more importantly, with a ∼1.5-fold increase in enzyme activity compared with that of wild type (WT). This work thus sets a solid foundation for future applications of LTA in biocatalytic synthesis of industrially important functional HAAs.

## RESULTS AND DISCUSSION

### Screening and characterization of LTAs from marine microbes

To search for LTAs with potential enzymatic activities, we used the synthetic reaction of l-*threo*-MPTS from 4-MTB and glycine as a model reaction (Scheme 1). A neighbor-joining phylogenetic tree was constructed for several LTAs from different sources according to their sequence similarity, which included 18 marine and 6 terrigenous microbial LTAs, among which three LTAs with available crystal structures were also included.^26, 28^ Terrigenous-sourced LTAs were grouped in the same clade, whereas the ones from marine source could be divided into five clades (Figure 1A). For each marine clade, we selected one representative LTA namely *Ml*LTA (from *Mameliella* sp.), *Sp*LTA (from *Sulfitobacter pontiacus*), *Al*LTA (from *Actibacterium lipolyticum*), *Nm*LTA, and *Rp*LTA (from *Rhodopirellula* sp.), and overexpressed these in heterologous *Escherichia coli* (*E. coli*) using pET22b with a his-tag. For comparison, three terrigenous LTAs with available crystal structures, i.e., *Aj*LTA (from *Aeromonas jandaei* DK-39), *Pp*LTA (from *Pseudomonas putida*), and *Ec*LTA (from *E. coli*), were overexpressed in *E. coli* (Figure S2).^26, 28^ We then examined the enzymatic activities of these LTAs against the model reaction using high performance liquid chromatography (HPLC) assays.^38^ N,N-dimethylformamide (DMF) was added as a co-solvent supporting the solubility of 4-MTB in the reaction (Table S3).

**Figure 1.**
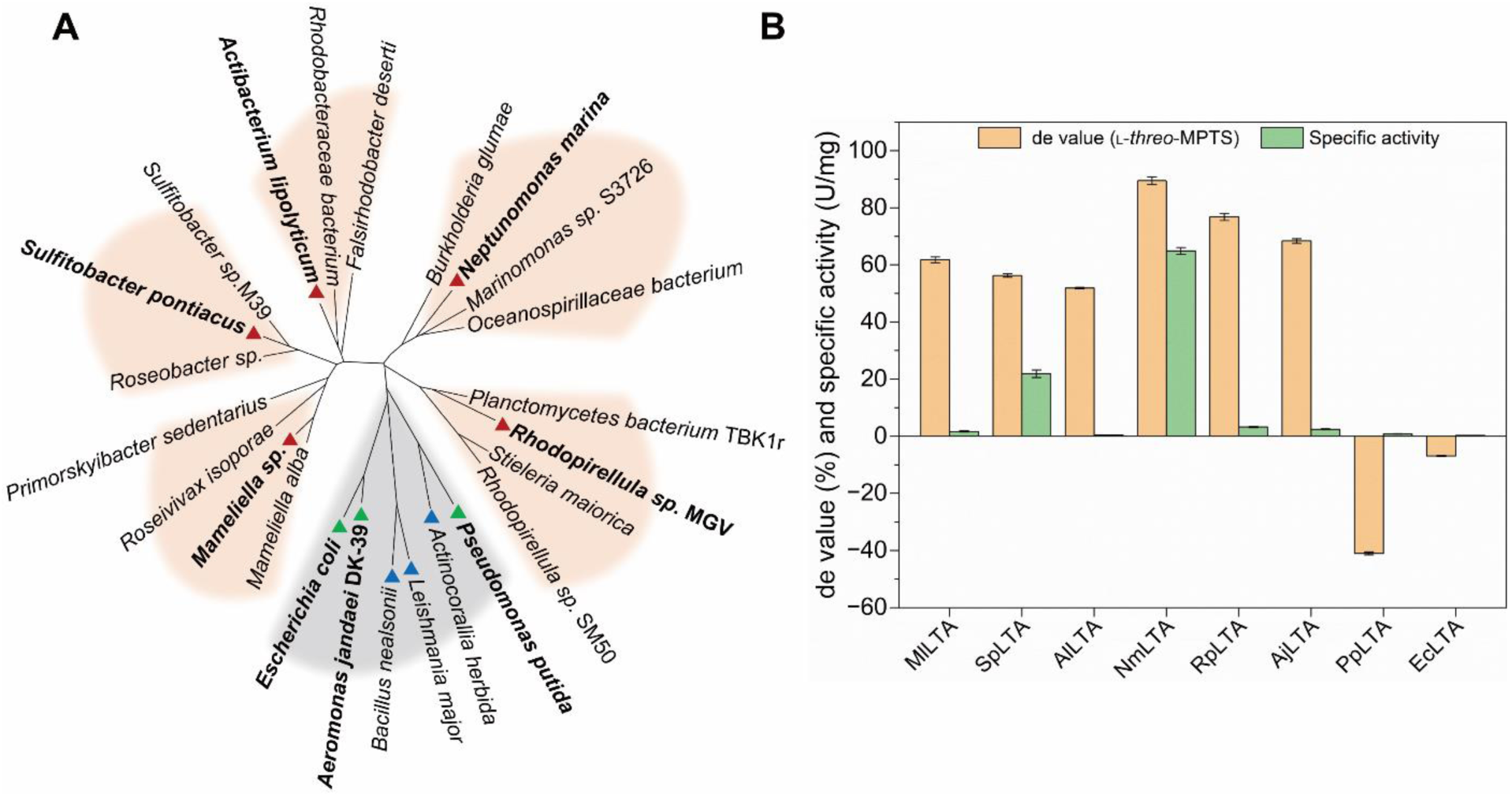
Mining and identification of LTAs with ideal activity and diastereoselectivity. (A) A simplified neighbor-joining phylogenetic tree showing the evolutionary relationship between characterized terrigenous LTAs and the newly discovered marine microbe LTAs. Marine and terrigenous microbe-sourced clades are colored in orange and gray background, respectively. Five candidate marine LTAs (e,g., *Ml*LTA, *Sp*LTA, *Al*LTA, *Nm*LTA, and *Rp*LTA) are labeled at the tree branches with red triangles. Three formerly characterized LTAs from *Bacillus nealsonii* (*Bn*LTA), *Leishmania major* (*Lm*LTA), and *Actinocorallia herbida* (*Ah*LTA) are labeled on the tree branches with blue triangles. Three LTAs (e.g., *Aj*LTA, *Pp*LTA, and *Ec*LTA) with available crystal structures are labeled at the tree branches with green triangles. All eight LTAs used in this study are highlighted in bold. (B) Specific activity and diastereoselectivity of LTAs for _L_-*threo*-MPTS. Reaction conditions: 100 mM aldehyde, 1 M glycine, 50 μM PLP, 10% DMF, and 20 μg/mL purified LTAs in 1 mL of 50 mM HEPES-NaOH buffer (pH 8.0) at 25 °C and 250 rpm within 30 min.

Most of the tested LTAs favored the *threo*-configuration product, whereas only *Pp*LTA and *Ec*LTA preferentially produced the *erythro*-configuration (Figure 1B and Table S1). Notably, all marine LTAs had a de value of > 50% for l-*threo*-MPTS. Among these, *Nm*LTA exhibited the highest de value of 89.5% and an efficient enzymatic activity (64.8 U/mg) that was ∼1.4-fold higher than the highest native LTA reported so far (Figure 1B and Table 1).^33^ Further analyses showed that *Nm*LTA had a maximum enzymatic activity at 30 ℃ and pH 8.0, and the efficiency could be further improved by increasing the PLP concentration (Figure S4). For thermal stability, *Nm*LTA remained highly active when incubated at 25 ℃ for over 140 min and remained 60% of its maximal at 50 ℃ over 60 min (Figure S5). Kinetic analyses for *Nm*LTA determined the respective *k*_cat_ and *K*_m_ values of 190.4 s^−1^ and 4.5 mM for l-*threo*-MTPS and 805.7 s^−1^ and 12.3 mM for the natural substrate l-threonine (Tables S2 and S7). Thus, *Nm*LTA is a promising enzymatic tool for establishing C–C bond owning to the ideal activity, which established a good foundation for continued attempts to improve the diastereoselectivity and activity.

**Table 1.** Comparison of the de values and specific activities for the synthesis of _L_-threo-MPTS by NmLTA and other identified LTAs.

| Organism | Enzyme | WT or variant | de value | Specific Activity (U/mg) |
| --- | --- | --- | --- | --- |
| *Pseudomonas* sp.<sup>12, 14</sup> | PsLTA | WT | 53.0% | - |
|  |  | V200I | 71.0% | - |
| *Bacillus nealsonii*<sup>30, 31</sup> | BnLTA | WT | 30.3% | 2.9 |
|  |  | Y31H/N305R | 93.1% | 1.2 |
|  |  | Y8H/Y31H/I143R/N305R | > 99% | - |
| *Actinocorallia herbida*<sup>19, 23</sup> | AhLTA | WT | 61.0% | 2.5 |
|  |  | Y314R | 81.0% | - |
|  |  | N16A/E98S/Y314R | 93.7% | - |
| *Leishmania major*<sup>32</sup> | LmLTA | WT | 26.8% | 17.4 |
|  |  | Y45H/Y319F/V321W | 96.3% | 12.1 |
| *Cellulosilyticum* sp.<sup>33</sup> | CsLTA | WT | 37.0% | 46.3 |
|  |  | Y8H/V143R/H305L | > 99% | 9.3 |
| *Neptunomonas marina* | NmLTA | WT | 89.5% | 64.8 |
|  |  | N18S/Q39R/Y319L | > 99% | 95.7 |

### Structural features of PLP-bound *Nm*LTA complex

To investigate the structural features of *Nm*LTA, we used X-crystallography to obtain the crystal structure of *Nm*LTA complexed with the PLP cofactor. After screening 1,600 crystallization conditions, the complex structure was resolved at a resolution of 2.8 Å (PDB ID: 7YVR) (Figures 2A, S6, and Table S4), which prompted us to dig deeper into the catalytic mechanism of the C_β_ stereoselectivity. *Nm*LTA belongs to the canonical PLP-dependent fold-type I enzyme family and has a homo-tetramer (Figure 2A).^11^ The functional group of PLP forms a Schiff base (internal aldimine) with the highly conserved K214 residue and directly interacts with *Nm*LTA residues at the monomer interfaces (inserted in Figure 2A). PLP can form π–π stacking interaction with the imidazole group of H95 and establish several hydrogen bonds (HBs) with D181 and R184 via the hydroxyl and imine groups. Moreover, G70, S71, and K239 stabilized the PLP phosphate group. Comparisons of the active-site residues of *Nm*LTA with that of other LTAs (*Pp*LTA, *Ec*LTA, and *Aj*LTA)^26, 28^ indicate that these residues are structurally highly conserved, highlighting their critical roles in substrate binding and catalysis (Figures 2B and S3).

**Figure 2.**
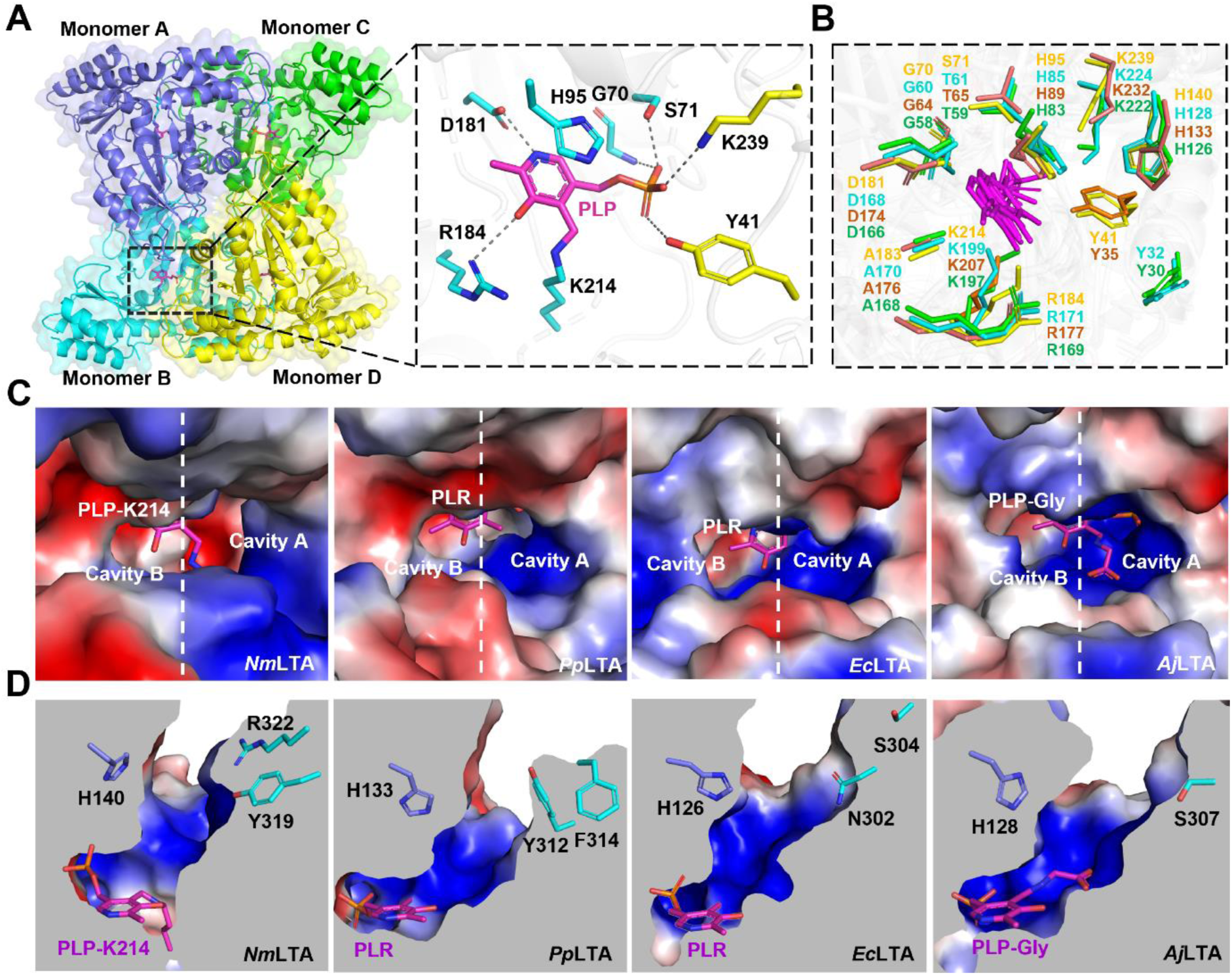
Crystal structure of *Nm*LTA and comparisons with those of other LTAs. (A) Overall structure of PLP-bound WT *Nm*LTA complex. The asymmetric unit of protein contains four monomers (A–D) in the P121 space group (left panel). The PLP cofactor (magenta sticks) is attached to the highly conserved K214 via forming a Schiff base and interacts with the LTA residues in two adjacent monomers (cyan and yellow). Key hydrogen bonds formed between PLP and the surrounding residues are highlighted in gray dashed lines. (B) Structural comparisons of active site residues from different LTAs, including *Nm*LTA (yellow), *Ec*LTA (green), *Aj*LTA (cyan), and *Pp*LTA (deep salmon). The PLP cofactors from all sources are shown as magenta sticks. Crystal structures of *Nm*LTA, *Pp*LTA, *Ec*LTA, and *Aj*LTA are derived from the Protein Data Bank identities: 7YVR, 5VYE, 4LNJ, and 3WGB, respectively. (C) Electrostatic potential surface of the substrate channel in four LTAs, colored from red (acidic) to blue (basic). PLP, PLR (4’-deoxypyridoxine phosphate), PLP-Gly, and PLP-K214 (LLP) are shown as magenta sticks. (D) Comparisons of the residues along the substrate channel in four LTAs.

We then compared the potential substrate binding pockets for different LTAs, as two large potential binding cavities (A and B) can be observed in all LTAs (Figure 2C). Analyses of the surface electrostatic potential for both cavities revealed that *Nm*LTA exhibits similar electrostatic properties to *Pp*LTA, with the former displaying more positive charges in cavity A than the latter because of the presence of R322. In sharp contrast, *Ec*LTA and *Aj*LTA demonstrated quite different electrostatic surfaces compared with those of *Nm*LTA. Additional structural analyses of cavity A show that *Nm*LTA contains a unique binding channel along which Y319 and R322 are located (Figure 2D), whereas either nonpolar (e.g., F314 for *Pp*LTA) or neutral residues (e.g., S304 for *Ec*LTA) are present in other LTAs (Figure 2D). Considering that the above four LTAs demonstrate distinct activities and diastereoselectivities for the synthesis of l-*threo*-MPTS (Figure 1B), we speculated that the varied residues and charge distributions in the LTA active site may dictate their different lead substrate specificities.

### Identification of potential *Nm*LTA residues responsible for diastereoselectivity control

After the binding of PLP to LTA, the catalysis proceeds first by binding of the α-amino acid (e.g., glycine) into the LTA active site, forming the PLP-Gly conjugate and displacing K214. According to the formerly proposed catalytic mechanism,^39^ the protonation of 4-MTB is likely promoted by a base-mediated water bridge. The activated PLP-Gly nucleophile will then attack the incoming aldehyde, leading to the C–C bond formation (Figure 3A). To scrutinize the structural features during the enzymatic aldol reaction, we first performed molecular docking for the intermediate PLP-Gly with *Nm*LTA. The energy-minimized *Nm*LTA-PLP-Gly complex revealed a similar binding mode for PLP to that observed in the *Nm*LTA-PLP complex. In addition, the carboxyl group of PLP-Gly could establish stable salt bridges with R184 and R329 and one HB with S16 (Figure 3B). This computational model was further supported by site-directed mutagenesis, which indicated that mutations of R184A and R329A could completely abolish the activity of *Nm*LTA, whereas the S16A mutant reduces the activity by approximately 80% (Figure 3C).

**Figure 3.**
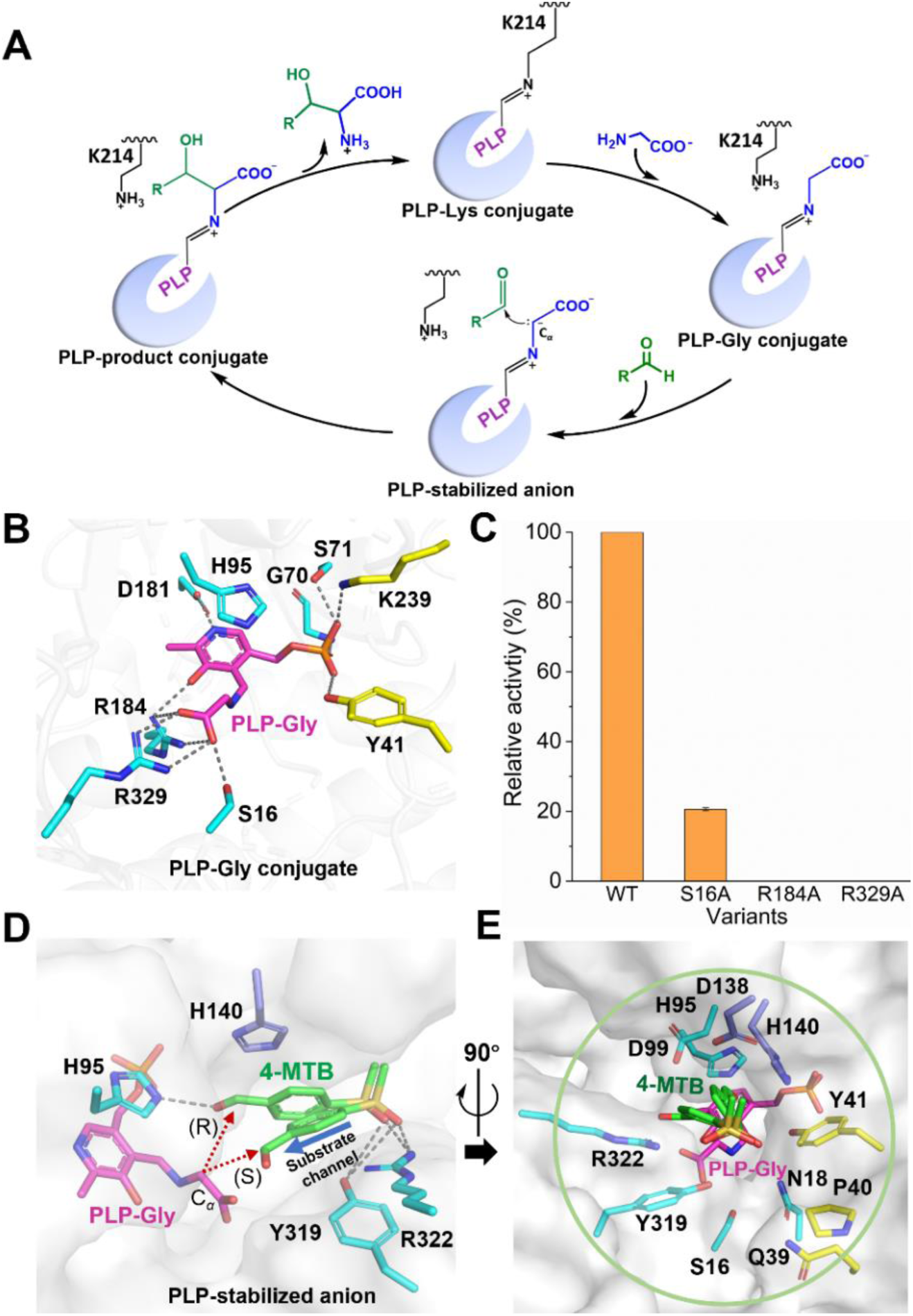
Computational modeling of substrate-bound *Nm*LTA complexes. (A) Schiff base exchange mechanism of *Nm*LTA. (B) Docking model of PLP-Gly-bound *Nm*LTA complex. Key HBs are highlighted in dashed lines. (C) Alanine scanning of residues surrounding the carboxyl group of PLP-Gly. (D) Docking models of ternary *Nm*LTA complexes where PLP-Gly and 4-MTB are shown as magenta and green sticks, respectively. Two docking conformations of 4-MTB are provided, corresponding to two configuration products (*R*/*S* configuration at C_β_-site). (E) Highlight of the residues within 6 Å surrounding the 4-MTB.

Using *Nm*LTA-PLP-Gly model, we docked the aldehyde substrate 4-MTB in the *Nm*LTA active sites. 4-MTB could potentially bind to *Nm*LTA in two different binding sites (namely, cavities A and B). Notably, in cavity B, 4-MTB formed few direct contacts with the *Nm*LTA residues, and moreover, the distance between the nucleophile PLP-Gly and the aldehyde group of 4-MTB was too distant (6.3 Å) to perform the reaction (Figure S7). Therefore, cavity B is an inappropriate binding site for 4-MTB. In comparison, in cavity A, 4-MTB fitted into a region flanked by R322, Y319, H140, and H95 where 4-MTB was sandwiched between H140 and Y319 and formed HBs with R322 and Y319 via the sulfonic-acid group (Figure 3D). Intriguingly, we observed two distinct orientations of the aldehyde group in 4-MTB, which probably gives rise to the two different aldol products (namely, *threo*- or *erythro*-form at the C_β_-site). The *threo*- configuration, which is the major product of *Nm*LTA, is located closer to H140 and forms one additional HB with H95 that is absent in the *erythro*-form, giving rise to a more favorable docking free energy (−4.9 and −3.8 kcal/mol for *threo*- and *erythro*-forms, respectively). Collectively, our findings suggest a “dual-conformation” mechanism for regulating the diastereoselectivity control of *Nm*LTA (Figure 3D).

These structural insights guided further attempts at enzyme engineering to improve the diastereoselectivity of *Nm*LTA. Previous studies reported that the activities of various LTA mutants were profoundly decreased, although with enhanced diastereoselectivity, such as for engineered *Bn*LTA, *Lm*LTA, and *Cs*LTA (Table 1).^30, 32, 33^ Here, to simultaneously improve the activity and diastereoselectivity, we first selected all the LTA residues located within 6 Å around the “dual-conformation” 4-MTB, which identified 11 residues, namely S16, N18, Q39, P40, Y41, H95, D99, D138, H140, Y319, and R322 (Figure 3E). Among these residues, the counterpart residues of residue H95 and D99 in other LTAs are known to play vital roles in substrate specificity and recognition, respectively.^23, 40^ Consistently, site-directed mutagenesis assays also proved that the H95A mutation could completely abolish the activity of *Nm*LTA (Figure S8). We also validated that P40, Y41, D138, and R322 are critical for specific activity or diastereoselectivity of *Nm*LTA and that the individual residue substitution would considerably impair either only the activity (i.e., D138) or both activity and diastereoselectivity (i.e., P40, Y41, D99, and R322) (Figure S8). Therefore, the remaining five residues (i.e., S16, N18, Q39, H140, and Y319) were selected for further systematic investigations.

### Site-saturation and combinatorial mutagenesis of selected *Nm*LTA residues

For each of above five chosen residues, we performed saturation mutagenesis using the NNK codon, and diastereoselectivity and activity screening were implemented using 4-MTB and glycine as substrates with a molecular ratio of 1:10, followed by HPLC analysis (Figures 4A and 4B). For each single mutation, both the de value and enzymatic activity were measured for l-*threo*-MPTS. For H140-site, all single substitutions produced a considerable decrease in the de value compared with that of the WT, and the H140L mutation had the lowest de value (−46.2%) (Figure 4A), indicating its essential role in regulating the diastereoselectivity. By contrast, we observed profoundly improved diastereoselectivity for several substitutions at the other four sites (Figure 4C). Specifically, we obtained a total of twelve *Nm*LTA variants with elevated de values ranging from 91.1% to 94.3% in the following order: S16A > S16G > Y319D > Y319L > Y319S > Q39R > N18S > Q39A > Q39L > Q39K > Q39H > N18T (Figure 4C). For enzymatic activity, except for Y319D and mutations at S16 and N18, all other positive substitutions displayed essentially equal (e.g., Q39H, Q39L, Q39A, and Y319L) or even higher (e.g., Q39R, Q39K, and Y319S) activities compared with that of the WT (Figures 4B, 4C, and Table S6).

**Figure 4.**
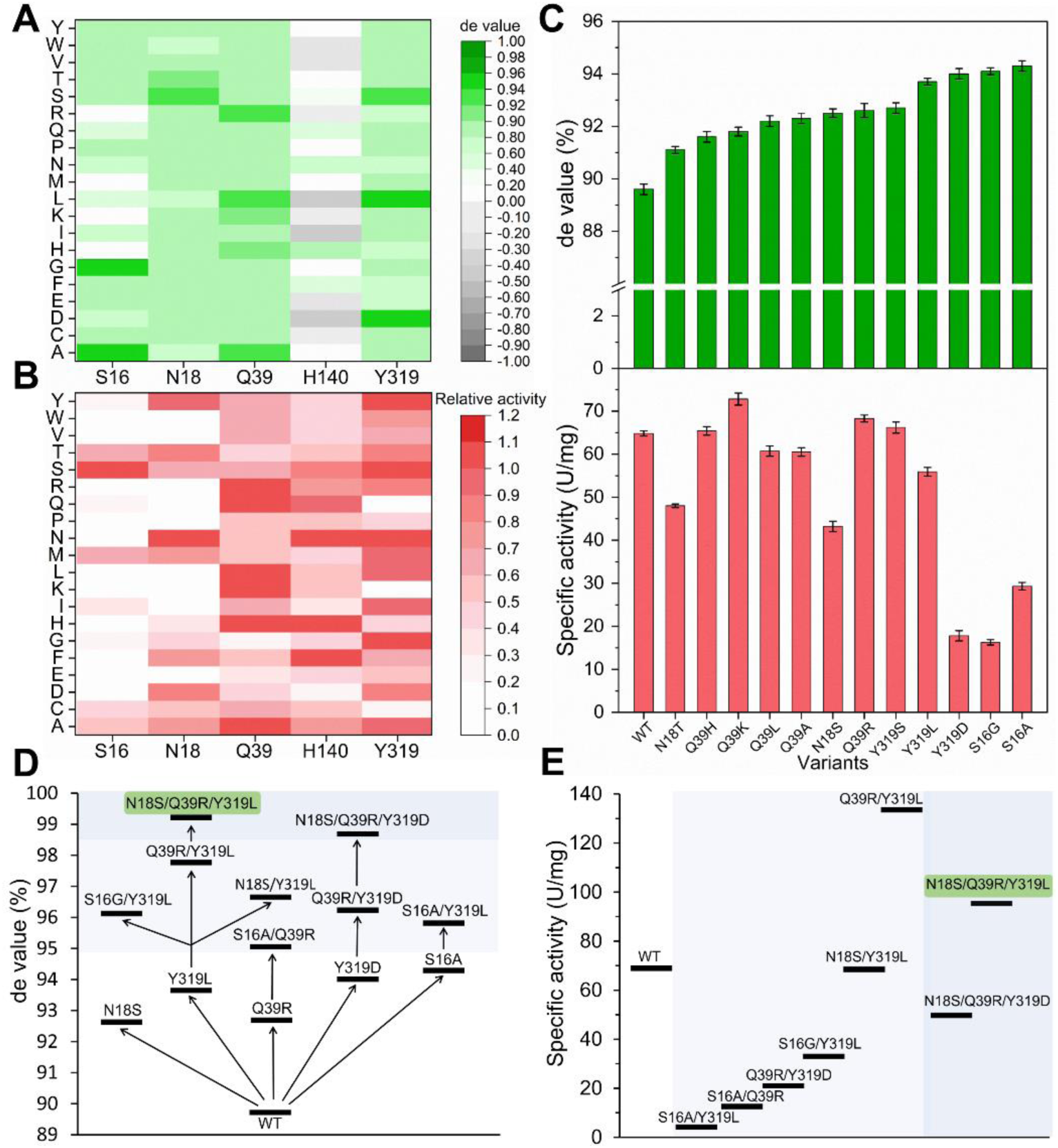
Saturation site-directed mutagenesis for evolving diastereoselectivity of *Nm*LTA. (A) Diastereoselectivities of *Nm*LTA mutants for the product _L_-*threo*-MPTS. (B) Relative activities of *Nm*LTA mutants for the product _L_-*threo*- MPTS and the catalytic activity of WT *Nm*LTA was defined as 100%. (C) The de value and specific activities of several positive mutants. (D) Development of combinatorial mutation strategy of LTAs with enhanced de value starting from the WT *Nm*LTA. (E) Specific activities of WT and different *Nm*LTA variants. Reaction conditions: 100 mM 4-MTB, 1 M glycine, 50 μM PLP, 10% DMF, and 20 μg/mL purified LTAs in 1 mL of 50 mM HEPES buffer (pH 8.0) at 25 °C and 250 rpm within 30 min.

To further enhance the diastereoselectivity and activity of *Nm*LTA, we conducted combinatorial mutagenesis based on the above twelve single mutants. Screening of the double-mutant libraries produced six candidates that exhibited enhanced diastereoselectivity, including S16G/Y319L, Q39R/Y319L, N18S/Y319L, S16A/Q39R, Q39R/Y319D, and S16A/Y319L (Figure 4D). Intriguingly, Q39R/Y319L showed the highest diastereoselectivity among all double-mutant variants (de value of 97.6%), and critically, it also had a 2-fold higher specific activity (133.0 U/mg) than that of the WT (64.8 U/mg) (Figures 4D, 4E, and Table S6). In addition, the *k*_cat_ and substrate performance values for the Q39R/Y319L mutant increased to 400.3 s^−1^ and 160.1, respectively, which were ∼2 and 6-fold higher than that of the WT (Table S7). We then performed a third round of screening based on these double mutants and identified two triple mutants, namely N18S/Q39R/Y319L (SRL) and N18S/Q39R/Y319D (SRD), with further improved the de values of 99.3% and 98.5%, respectively (Figure 4D and Table S6). Notably, the SRL variant had a specific activity of 95.7 U/mg and a *k*_cat_/*K*_m_ of 78.9 s^−1^mM^-1^, which were ∼1.5-fold and 1.9-fold higher than those of the WT, respectively. Moreover, the SRL variant displayed a ∼10.7-fold higher substrate performance than that of the WT (297.5 vs. 27.8, respectively) (Figure 4E and Table S7). Altogether, the SRL variant of *Nm*LTA not only showed ideal diastereoselectivity for l-*threo*-MPTS but also displayed improved enzymatic activities compared with that of the WT.

### Dual-conformation regulation for the diastereoselectivity control of *Nm*LTA

As described above, our computational studies indicated that 4-MTB could potentially bind at the *Nm*LTA cavity A in two distinct orientations, clamped by the sidechains of H140 and Y319 sidechains. Thus, the activated PLP-Gly could potentially attack the carbonyl group of 4-MTB from two possible directions, resulting in the *threo*- or *erythro*-product (Figure 5D and Figure S9). We therefore proposed a “dual-conformation” regulation for the diastereoselectivity control of *Nm*LTA whereby H140 and Y319 play key roles in fine-tuning the substrate conformation. The proposed mechanism was supported by site-directed mutagenesis, which indicated that substitutions of H140 with any other residues could profoundly decrease the de value for the *threo*- configuration product and most of these mutants underwent diastereoselective inversion, whereas mutations of Y319 to Leu, Ser, or Asp had a reverse effect. In comparison, we performed a saturation mutation of V155 that is located in cavity B of *Nm*LTA. Notably, the mutation of the counterpart residue V155 in *Bn*LTA (I143R) and *Cs*LTA (V143R) enhanced de values (Table 1).^31, 33^ By contrast, none of the variants at the V155 site exerted any considerable influence on the diastereoselectivity for the synthesis of l-*threo*-MPTS (Figure S10), supporting our conclusion that cavity B does not participate in regulating the substrate diastereoselectivity. This work therefore rules out the formerly proposes “two pathways” mechanism as observed in other LTA,^30, 31, 33^ and proposes a novel diastereoselectivity mechanism for *Nm*LTA. Interestingly, a similar mechanism was also observed in a pyruvate-dependent aldolase, where the pyruvate enolate could attack the *si* or *re* face of the substrate succinic semialdehyde to yield racemic products.^41^

**Figure 5.**
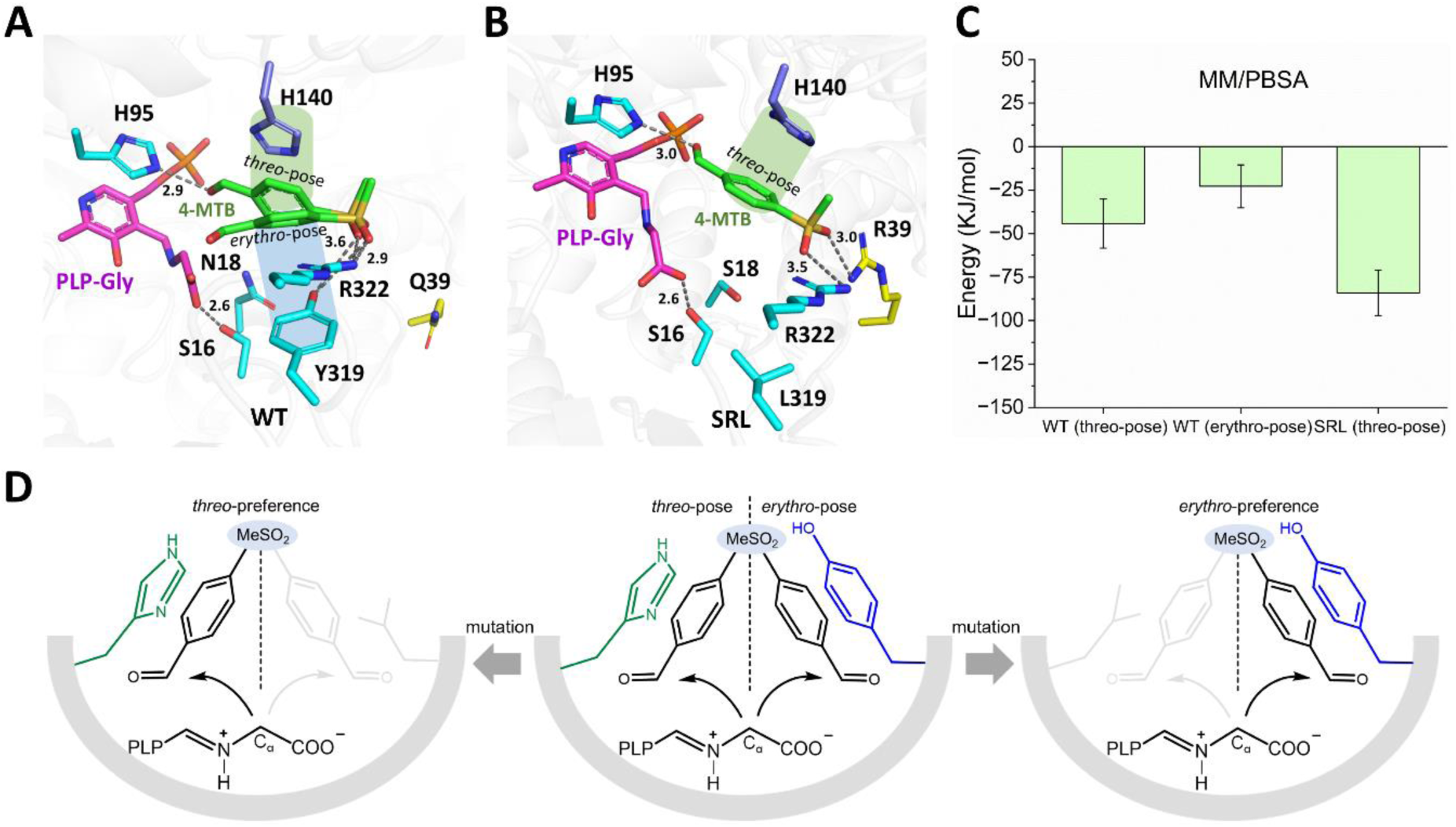
The “dual-conformation” regulation in *Nm*LTA. (A) Conformation of 4-MTB in WT *Nm*LTA. Residues Q39, H140, Y319, and R322 line the active site channel (labeled by purple, cyan, and yellow, respectively). Gray dashed lines represent hydrogen bond interactions. (B) The conformation of 4-MTB in the SRL variant. The grey dashed lines represent hydrogen bond interactions. (C) Binding free-energy calculation of the WT *Nm*LTA and the SRL variant using the MM/PBSA algorithm during the initial 1 ns. (D) Dual-conformation regulation for the diastereoselectivity control of *Nm*LTA.

We next compared the structural difference between the WT *Nm*LTA and the SRL variant by constructing an SRL model based on the WT crystal structure. Structural inspections revealed that the SRL variant can profoundly alter the binding mode of the incoming aldehyde in the active site. Specifically, the Y319L substitution weakened the packing interactions between the aromatic ring of the aldehyde and Y319 as observed in the WT, which subsequently pushed the substrate toward H140, leading to the preferred formation of the *threo*-product (Figures 5A, 5B, and 5D). This finding was consistent with the above observations that all the single mutants at H140 yielded reduced de value for l-*threo*-MPTS (Figures 4A and 5D). In addition, the Q39R mutation provided an additional anchor point to stabilize the sulfonic acid tail of 4-MTB together with R322, which likely increased enzymatic activity (Figures 5A and 5B). Moreover, the N18S mutation likely imposed unsubstantial structural perturbations on PLP-Gly by altering the HB interaction with Y41, slightly improving diastereoselectivity (Figure S11). Consistently, our binding free-energy calculations showed that the SRL variant of *Nm*LTA demonstrated stronger binding affinity between the aldehyde and protein compared with that of the WT (Figure 5C), and that this was mainly caused by the additional salt bridge interaction between R39 and the sulfonic-acid group of 4-MTB. Notably, previous studies of other LTAs had identified counterpart residues of *Nm*LTA H140, Y319, and N18 (e.g., H136 and Y314 in *Ah*LTA, H142 and Y319 in *Lm*LTA, Y8H in *Bn*LTA and N16A in *Ah*LTA), suggesting that these residues may serve as universal sites for diastereoselectivity control (Figure S12).^31, 32^ The Q39 position, however, has not yet been examined for other LTAs, therefore serving as a potential engineering site for other LTAs.

### Catalytic performance of the SRL variant in preparative gram-scale l-*threo*-MPTS

To evaluate the performance of *Nm*LTA for the potential industrial synthesis of l-*threo*-MPTS, we first examined the thermal stability of the SRL variant. The results showed that the SRL variant maintained > 98% activity after heating at 25 ℃ for 140 min and still retained 70% of the activity at 50 ℃ for 60 min, demonstrating improved thermostability over that of the WT (Figure S5). We then evaluated the gram-scale synthesis of l-*threo*-MPTS in a 1 L scale reactor by including 100 mM 4-MTB and 1 M glycine with 1,400 U WT *Nm*LTA or the SRL variant. Measurement of the de value and yield of l-*threo*-MPTS indicated that the SRL variant always maintained strict diastereoselectivity with increased conversion rate, whereas for the WT, the de value slightly decreased as the conversion rate increased. For the SRL variant, the yield of l-*threo*-MPTS was 21.0 g/L, with a de value of 99.3% and space-time yield of 9.0 g L^−1^h^−1^. Thus, the SRL variant greatly improved the catalytic performance of *Nm*LTA, displaying much stricter diastereoselectivity compared with that of the WT (Figure 6). Therefore, the SRL variant could potentially serve as a robust C–C bond synthetic tool for industrial synthesis of HAAs on a preparative scale.

**Figure 6.**
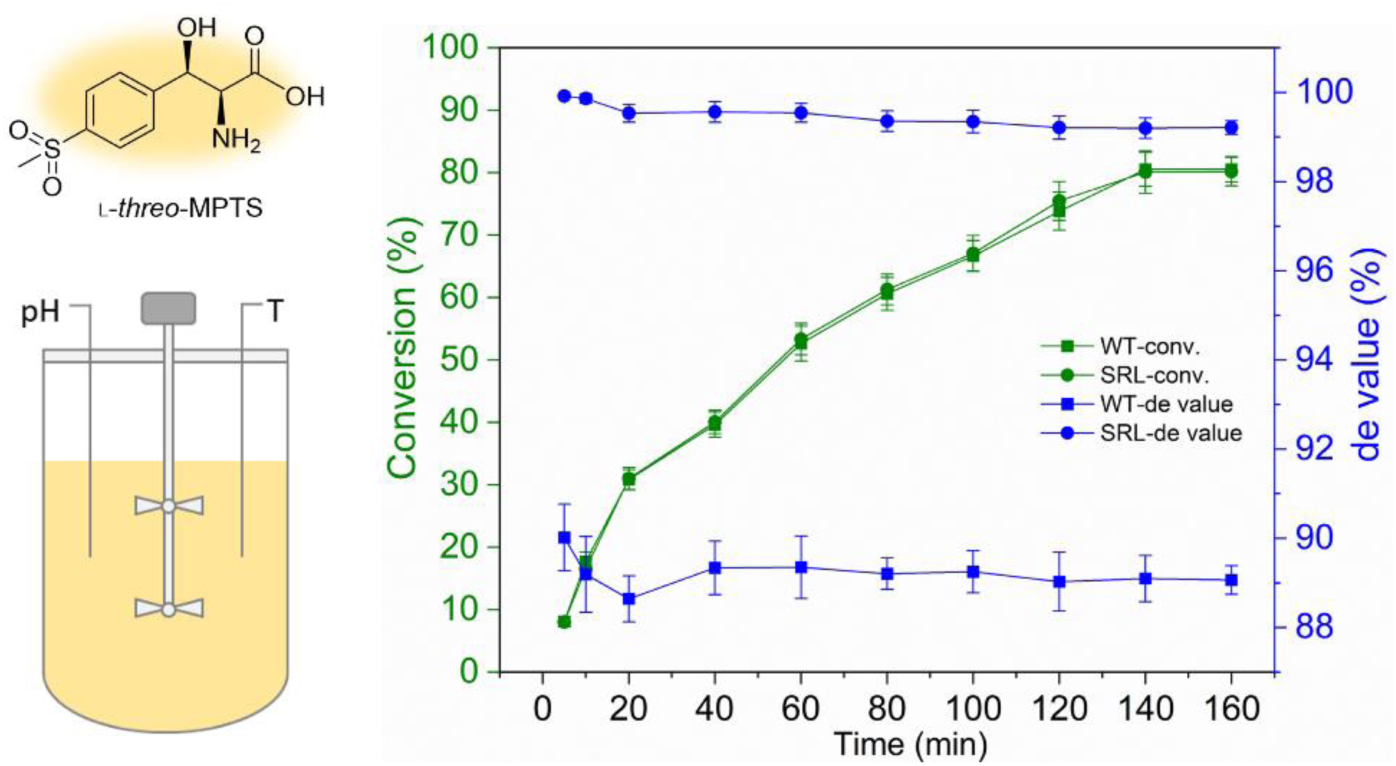
Catalytic performances of WT *Nm*LTA and the SRL variant for the synthesis of _L_-*threo*-MPTS. Reaction conditions: 100 mM 4-MTB, 1 M glycine, 50 μM PLP, 10% DMF, and 1400 U LTAs in 1 L of 50 mM HEPES buffer (pH 8.0) at 25 °C and 250 rpm within 160 min.

### Substrate scope of the evolved *Nm*LTA variant

To further examine the substrate scope of the WT *Nm*LTA and the SRL variant, we investigated the enzymatic activities and diastereoselectivities of LTAs toward a representative library of benzaldehyde derivatives. The WT *Nm*LTA and the SRL variant could recognize and catalyze all tested substrates except for the 4-OH substituted benzaldehyde, with the conversion rates varying between 58% and 80% (Figure 7). As listed in Figure 7, all *para*-substituted substrates had increased de values except for 4-F (de value > 80%), in particular, for 4a, the de value could reach up to 85.7%. For the *meta*-site, only 3-Br-substituted substrate improved the de value, whereas substitution with 2-Cl or 2-NO_2_ at *ortho*-site increased the de values. Moreover, two tested disubstituted substrates, 3,4-CH_3_ and 3,5- CH_3_, undermined the enzymatic performance, which is likely due to the large steric hindrances caused by the substituents. For substrate benzaldehyde, the SRL variant shows a considerable increase in the de value (12a). Altogether, the SRL variant shows a broad substrate scope for the synthesis of diverse HAAs, which are important synthetic precursors for bioactive compounds and therapeutic drugs (e.g., 5a for chloramphenicol). Notably, different substrates with varied substituents could apparently impose considerable influences on the enzymatic activities and diastereoselectivity control of the SRL variant, likely caused by the perturbed binding mode of the substrates. Despite that, the SRL variant appeared more tolerant toward the *para*-substituted substrates by electronegative or neutral groups (except for 4-F).

**Figure 7.**
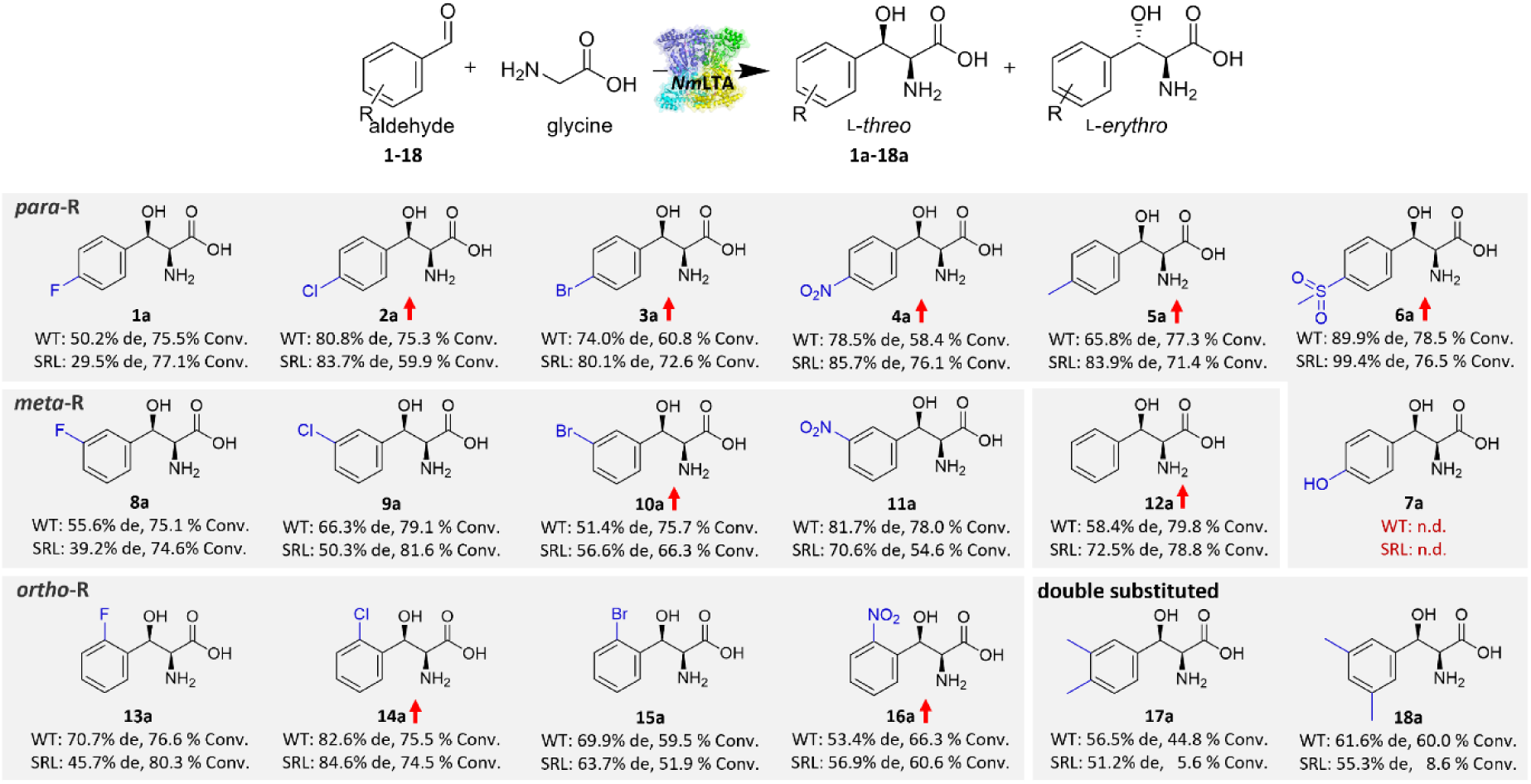
Diastereoselectivities and conversions of WT *Nm*LTA and the SRL variant for the synthesis of HAA 1a– 18a. Reaction conditions: 100 mM aldehyde, 1 M glycine, 50 μM PLP, 10% DMF, and 1.5 U LTAs in 1 mL of 50 mM HEPES buffer (pH 8.0) at 25 °C and 250 rpm within 120 min. See Figures S13–S29 for all liquid chromatograms and mass spectrograms.

## CONCLUSION

Herein, we report a robust LTA from marine *N. marine* (*Nm*LTA) that established the C–C bond with high enzymatic activity and strict diastereoselectivity control of the C_β_-site. X-ray crystallography, computational modeling, and site-directed mutagenesis allowed us to design a triple mutant of *Nm*LTA, namely the SRL variant, that exhibits > 99% de value for l-*threo*-MPTS, and importantly, demonstrates ∼1.5-fold increased enzymatic activity (95.7 U/mg) compared with that of WT to achieve simultaneous enhancement of activity and diastereoselectivity for the synthesis of l-*threo*-MPTS. Meanwhile, the SRL variant of *Nm*LTA exhibited the highest specific activity for synthesizing l-*threo*-MPTS among the currently reported LTAs (Table 1).^33^ Moreover, the preparative gram-scale aldol reaction of l-*threo*-MPTS catalyzed by the SRL variant produced a space-time yield of up to 9.0 g L^−1^h^−1^. Furthermore, the substrate spectrum assays indicated that the SRL variant could accept a wider range of aromatic aldehyde derivatives as substrates and show improved diastereoselectivity by tuning the *para*-site substituents. Our work identified the key structural motifs in *Nm*LTA responsible for regulating its activity and diastereoselectivity and proposed a novel molecular mechanism underlying the diastereoselectivity control of C–C bond formation, thereby accelerating the application of the aldol reaction using engineered LTAs for the biocatalytic preparation of industrially important functional HAAs.

## AUTHOR INFORMATION

### Supporting Information

Chemicals and Materials used in this study; Methods of gene mining, synthesis, expression, protein purification, site-directed mutagenesis, and screening of variant; Activity, kinetic, and substrate specificity assay; Crystallization and protein structure determination; Molecular docking, molecular dynamic simulations, and calculations of molecular mechanics Poisson-Boltzmann Surface Area; HPLC and MS Analysis; Primers and relative DNA sequences used in this study.

### Corresponding Author

* Yan Feng – State Key Laboratory of Microbial Metabolism, Joint International Research Laboratory of Metabolic & Developmental Sciences, School of Life Sciences and Biotechnology, Shanghai Jiao Tong University, 800 Dongchuan Road, Shanghai, China.

• Lin-Tai Da – Key Laboratory of Systems Biomedicine (Ministry of Education), Shanghai Center for Systems Biomedicine, Shanghai Jiao Tong University, 800 Dongchuan Road, Shanghai, China.

### Notes

The authors declare no competing financial interest.

## ACKNOWLEDGEMENTS

We thank staffs of beamlines BL18U1 and BL19U1 at Shanghai Synchrotron Radiation Facility for their assistance in diffraction data collection. We thank Prof. Juan Lin (College of Chemical Engineering, Fuzhou University, China) for providing the product standard.

