## Supplementary material for "Discovery and Engineering of the l-Threonine Aldolase from *Neptunomonas Marine* for Efficient Synthesis of β-Hydroxy-α-Amino Acids via C–C Formation": Supporting Information.pdf

### 1.1 Chemicals and Materials

4-methylsulphonylbenzaldehyde (4-MTB) and all benzaldehyde derivatives were purchased from Adamas Reagent. Isopropyl β-d-1-thiogalactopyranoside (IPTG), nicotinamide adenine dinucleotide (NADH), threonine, and glycine were purchased from Sangon Biotech. Alcohol dehydrogenase (ADH) was purchased from Sigma-Aldrich. Pyridoxal-5'phosphate (PLP) was obtained from TCI. Acetonitrile was purchased from Merck. All restriction enzymes were purchased from Thermo Fisher Scientific. High-fidelity DNA polymerase (PrimeSTAR® Max) was purchased from Takara. Other reagents were purchased from Sinopharm Chemical Reagent.

### 1.2 Experimental Section

#### 1.2.1 Gene Mining, Synthesis, Expression, and Protein Purification

The previously reported protein *Pp*LTA was used as the query sequence using BLASTP algorithm in GenBank non-redundant protein sequence database to identify potential LTAs[1]. Bioinformatics filters (identity > 40% and < 80%; containing key motifs of *Pp*LTA; host of marine microorganisms) were used to recruit potential protein sequences. The results of the homologous sequences were then processed by phylogenetic analysis using MEGA 11 with neighbor-joining algorithms[2]. All LTAs used in this study were listed in Table S1. Codon-optimized versions of the genes for candidate LTAs were performed from GenScript (Nanjing, China) and cloned into pET22b vector between NdeI and XhoI sites. *E. coli* BL21(DE3) was used for heterologous overexpression of LTA proteins and the mutated proteins. Site-directed mutagenesis for the creation of *Nm*LTA variants were performed by Tsingke Biotech (Beijing, China). Cultivation was performed in Luria-Bertani (LB) medium with ampicillin (100 μg/mL). Cultures were initially incubated at 37 °C with shaking at 220 rpm. At an $OD_{600}$ of between 0.6 and 0.9, IPTG was added to a final concentration of 0.5 mM to induce the expression of LTAs. Incubation was continued at 25 °C and 220 rpm for 8-12 h, the cells were harvested by centrifugation and resuspended in HEPES buffer (50 mM, pH 8.0) containing 300 mM NaCl and 50 mM imidazole. Cells were disrupted by ultrasonication and the soluble protein was purified by Ni-affinity chromatography. Purified proteins were examined by SDS-PAGE. The protein concentration was determined using the Bradford assay against BSA as a concentration standard. We used a centrifugal concentrator (Millipore) with a molecular weight cutoff of 10 kDa to remove the imidazole and the purified proteins were stored at −80 °C until used. Approximately 60 mg of purified LTAs could be obtained from a 1 L LB culture.

#### 1.2.2 Activity, Kinetic, and Substrate Specificity Assay

To determine the activity of LTA for natural substrate, a typical reaction mixture contained 50 mM L-threonine, 50 μM PLP, 2 mM NADH and 10 U ADH in HEPES buffer (50 mM, pH 8.0) at 25 °C, determination of NADH concentration monitored at 340 nm ($\varepsilon = 6.22\ mM^{-1}cm^{-1}$). A boiled enzyme sample was used as a negative control. One unit (U) was defined as the amount of enzyme reducing 1 μmol of NADH in one minute. The kinetic parameters ($K_m$ and $k_{cat}$) were measured with variable substrate concentrations between 0.5 and 50 mM at 25 °C and calculated according to Michaelis-Menten plot. To determine the activity of LTA for unnatural substrate, a typical reaction mixture contained 100 mM aldehyde, 1 M glycine, 50 μM PLP, and 10% DMF in HEPES buffer (50 mM, pH 8.0) at 25 °C, 250 rpm. A boiled enzyme sample was used as a negative control. One unit (U) of aldol reaction activity was defined as the amount of enzyme catalyzing the synthesis of 1 μmol of HAAs in one minute. The kinetic parameters ($K_m$ and $k_{cat}$) were measured with variable substrate concentration between 0.1 and 40 mM at 25 °C and calculated according to Michaelis-Menten plot.

#### 1.2.3 Crystallization and Structure Determination

Purified *Nm*LTA was subjected to crystallization trials using a range of commercially available screens in 96-well sitting-drop format in which each drop consisted of 0.2 μL protein and 0.2 μL of precipitant reservoir solution. Finally, long strip diffraction quality crystals were obtained in the presence of 100 mM sodium acetate (pH 4.75), 23% (v/v) 2-methyl-2,4-pentanediol. Crystals were cryoprotected using a reservoir solution containing 25% (v/v) glycerol. The X-ray data were collected on BL18U1 beamline at the Shanghai Synchrotron Radiation Facility (SSRF). All diffraction data were indexed, integrated, and scaled using XDS-GUI[3]. The structure was solved by molecular replacement methods using the crystal structure of phenylserine aldolase (PDB ID: 1V72) as the template. Subsequently, the structure was refined using Phenix[4] and inspected with the program COOT[5, 6]. The final model was evaluated using the MolProbity program[5]. Data collection, processing, and refinement statistics were summarized in Table S4. The *Nm*LTA coordinates and structure factors have been deposited in the PDB under accession number 7YVR. Graphic representations of structures were generated in PyMoL[7].

#### 1.2.4 HPLC and MS Analysis

The products were used for detecting conversion and diastereoselectivity by high performance liquid chromatography (HPLC) with a UV detector after chiral derivatization with o-phthalaldehyde/N-acetyl-L-cysteine (OPA/NAC)[8]. The HPLC analysis were carried out on an Agilent1260 Series HPLC system and Eclopse SB-C18 column. The mobile phase was $KH_2PO_4$ solution (50 mM, pH 8.0)/$CH_3CN$ = 85/15 and the flow rate was 1 mL/min.

The column temperature was 30 °C and the detection wavelength was 340 nm. The reaction was quenched by adding 20 volumes of acetone and centrifuged to remove precipitated glycine. For MS analysis, the reaction was quenched by adding 10 volumes of methanol and centrifuged, and the supernatant was analyzed by Agilent 1290 Infinity II/6545 QTOF-LC/MS, ESI (+).

#### 1.2.5 Molecular Docking and Molecular Dynamic Simulations

We obtained binary and ternary complexes of LTA by sequentially docking PLP-Gly and 4-MTB to the crystal structure of *Nm*LTA using AutoDock 4.5 software[9]. For each substrate, 100 docking complexes were generated with default docking parameters, then the lowest docking complex was subject to the following molecular dynamic (MD) simulations using the Gromacs 2020 package[10]. The protein was described by the force field Amber ff14SB. To obtain the force field parameters of PLP-Gly and 4-MTB, the HF/6-31G* method of Gaussian 09 software package was used to calculate the electrostatic potential of small molecules, and then the Antechamber module from the Amber-Tools package was used to calculate the RESP charges[11]. Other parameters of small molecules were derived from GAFF. Each LTA complex was placed in a cubic box filled with SPC216 model water with a minimum distance of 12 Å from the boundary of the water box. Appropriated numbers of $Na^+$ and $Cl^-$ ions were added to neutralize the whole system and reach an ion-concentration of 0.15 M. Then, the 200-ps NVT equilibrium MD simulations were conducted by restraining all the heavy atoms of the system, followed by a 1-ns position-restrained equilibrium simulations at 298 K. The long-range electrostatic interactions were treated by Particle Mesh Ewald (PME) method[12]. The van der Waals and short-range electrostatic interactions were set to 12 Å. The LINCS algorithm was used to constrain all the bonds to correct the bond lengths[13]. The temperature was coupled using the velocity rescaling with a stochastic term[14].

#### 1.2.6 Calculations of Molecular Mechanics Poisson-Boltzmann Surface Area

We employed the Molecular Mechanics Poisson-Boltzmann Surface Area (MM/PBSA) method to calculate the binding free-energy between WT/SRL *Nm*LTA and the ligand[15]. We collected a total of 20 conformations for each LTA complex. Next, for each conformation, the binding free-energy was calculated using the MM/PBSA method implemented in gmx_mmpbsa[16]. Finally, the average and corresponding errors were calculated.

#### 1.2.7 Site-directed Mutagenesis and Screening of *Nm*LTA

We used whole-plasmid PCR (Polymerase Chain Reaction) to perform site-directed mutagenesis. The recombinant plasmid was used as the template and the primers used were shown in Table S5. The PCR amplification was carried out in 50 μL of a solution consisting of 25 uL DNA polymerase mix (2 × PrimeSTAR®, Takara), 2 μL F- and R-primer, 1 ng DNA template, and 20 μL dd$H_2O$. The PCR program was 94 ℃ for 3 min, 56 ℃ for 35 s, 72 ℃ for 1 min, a total of 30 cycles. The PCR products were recovered using the Clean-Up Kit (Axygen). And the final products were transformed to *E. coli* BL21 (DE3) competent cells, and cultured in the LB solid medium with 100 μg/mL ampicillin for 12-16 h. And then, these single colonies were cultured in LB liquid medium for sequencing. After verification by sequencing, the mutants were expressed and purified for *in vitro* reaction, and the products were detected by HPLC. Then the product concentration and de values were calculated by standard curves.

### 1.3 Supporting Figures

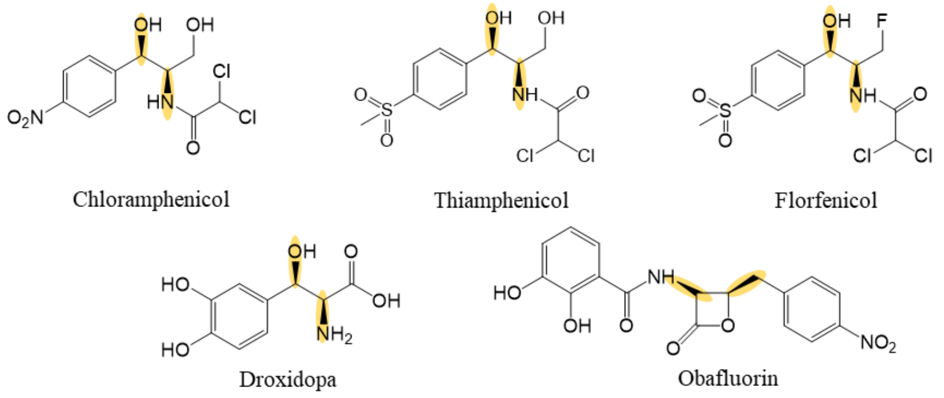

**Figure S1.** Complex β-hydroxy-α-amino acid building blocks in pharmaceuticals.

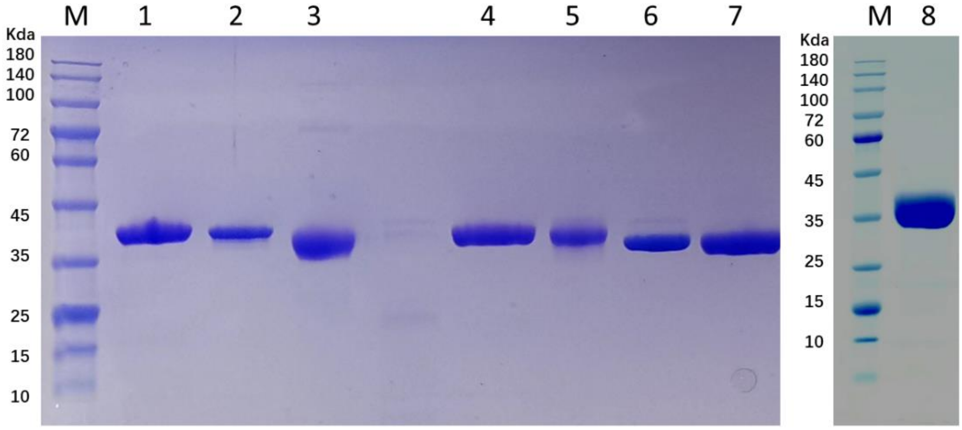

**Figure S2**. SDS-PAGE analysis of 8 purified LTAs. M: marker, lane 1-8: *Ml*LTA, *Al*LTA, *Nm*LTA, *Rp*LTA, *Sp*LTA, *Pp*LTA, *Ec*LTA, and *Aj*LTA. *Ml*LTA (*Mameliella* sp.), *Al*LTA (*Actibacterium lipolyticum*), *Nm*LTA (*Neptunomonas marina*), *Rp*LTA (*Rhodopirellula* sp. MGV), *Sp*LTA (*Sulfitobacter pontiacus*), *Pp*LTA (*Pseudomonas putida*), *Ec*LTA (*Escherichia coli*), and *Aj*LTA (*Aeromonas jandaei* DK-39).

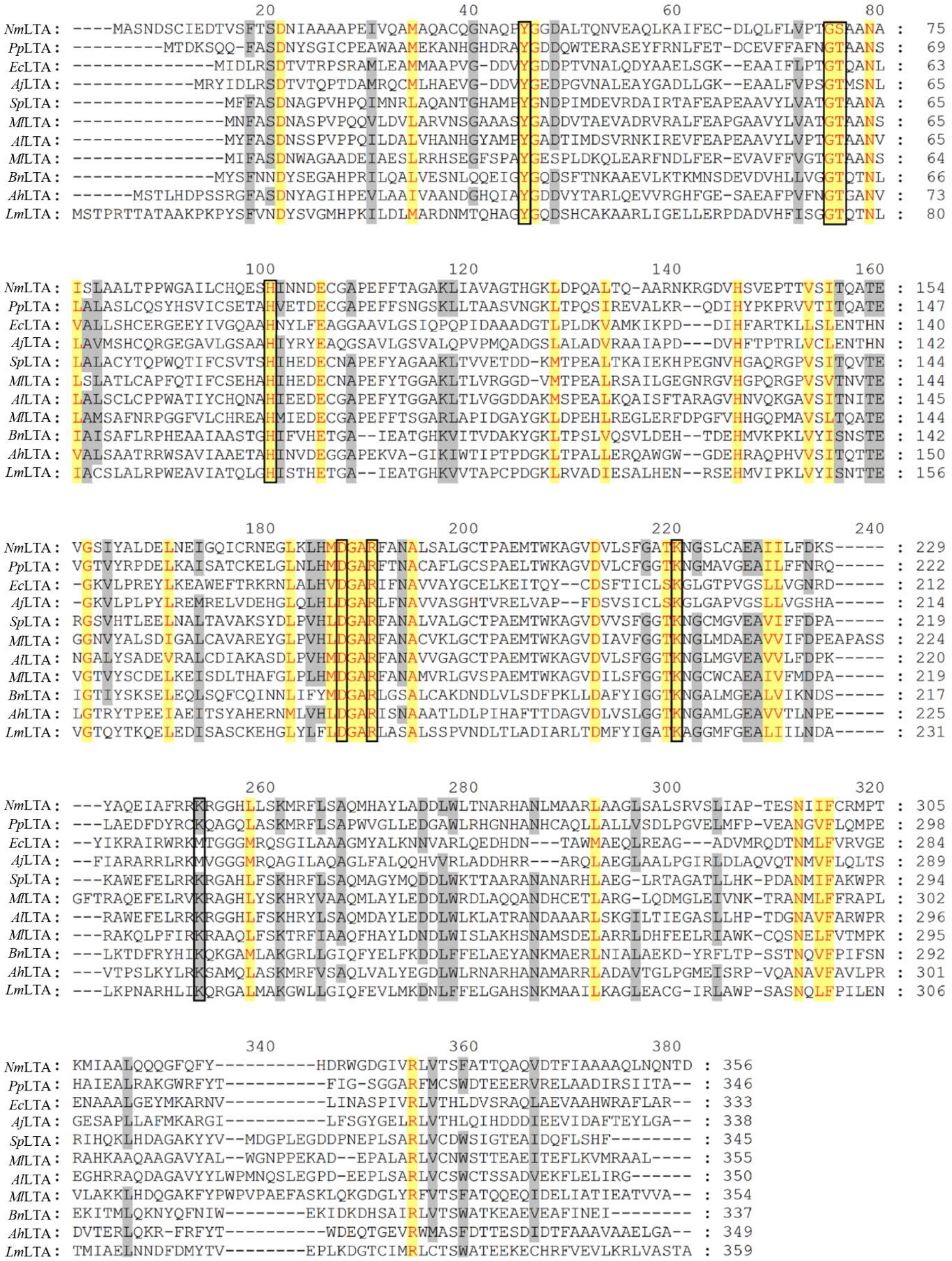

**Figure S3.** Amino acid sequences alignment of *Nm*LTA and other 10 LTAs. *Nm*LTA (*Neptunomonas marina*), *Pp*LTA (*Pseudomonas putida*), *Ec*LTA (*Escherichia coli*), *Aj*LTA (*Aeromonas jandaei* DK-39), *Sp*LTA (*Sulfitobacter pontiacus*), *Ml*LTA (*Mameliella* sp.), *Al*LTA (*Actibacterium lipolyticum*), *Rp*LTA (*Rhodopirellula* sp. MGV), *Bn*LTA (*Bacillus nealsonii*), *Ah*LTA (*Actinocorallia herbida*), and *Lm*LTA (*Leishmania major*). PLP binding residues are colored with black frames.

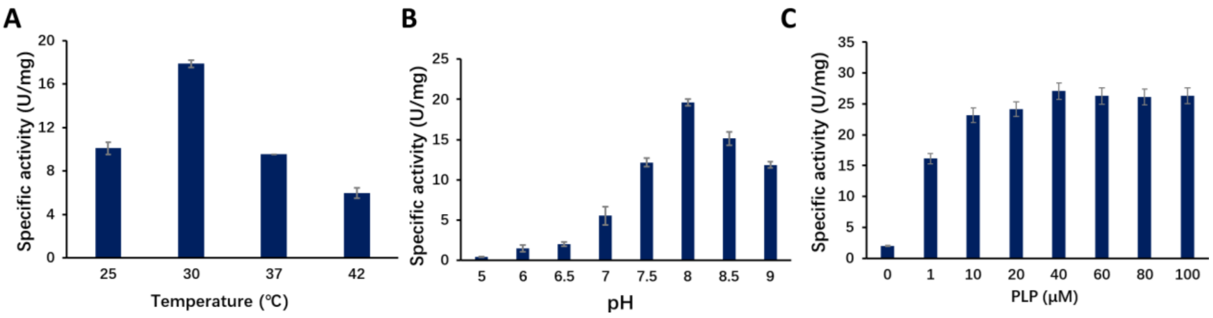

**Figure S4.** Enzymatic properties of *Nm*LTA for natural reaction. (A) Detection of the optimal reaction temperature of *Nm*LTA. (B) Detection of the optimal reaction pH of *Nm*LTA. (C) Effect of PLP concentration on the enzyme activity of *Nm*LTA. Reaction conditions: 50 mM L-threonine, 50 μM PLP, 2 mM NADH, 10 U ADH, and 10 μg purified LTAs in 50 mM HEPES buffer (pH 8.0) at 25 °C, determination of NADH concentration monitored at 340 nm. All experiments were conducted in triplicate.

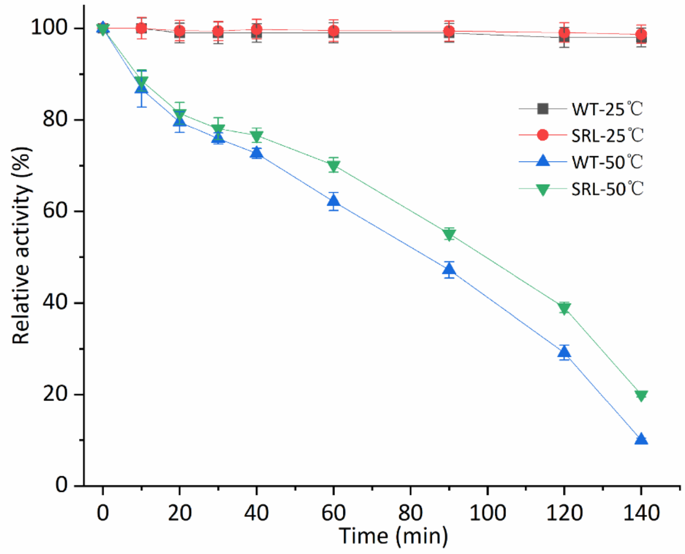

**Figure S5.** Temperature stability of *Nm*LTA and SRL variant after incubation of enzyme solution. Reaction conditions: 100 mM 4-MTB, 1 M glycine, 50 µM PLP, 10% DMF, and 20 µg purified LTAs in 1 mL of 50 mM HEPES buffer (pH 8.0) at 25 °C and 50 °C and 250 rpm within 30 min. All experiments were conducted in triplicate.

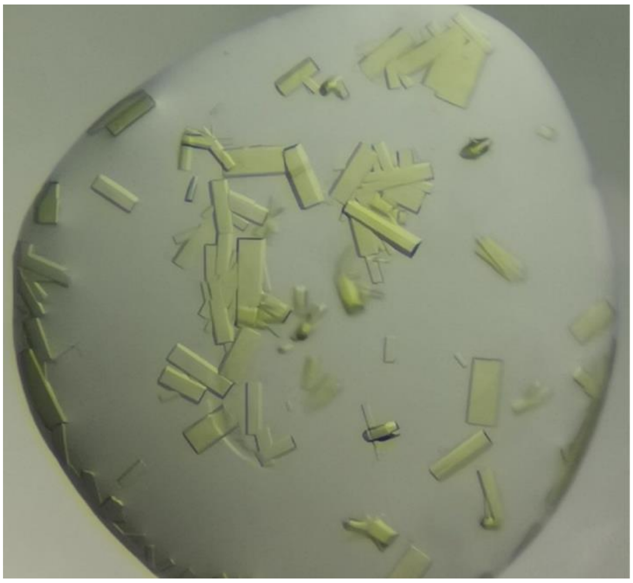

**Figure S6.** Crystals of WT *Nm*LTA.

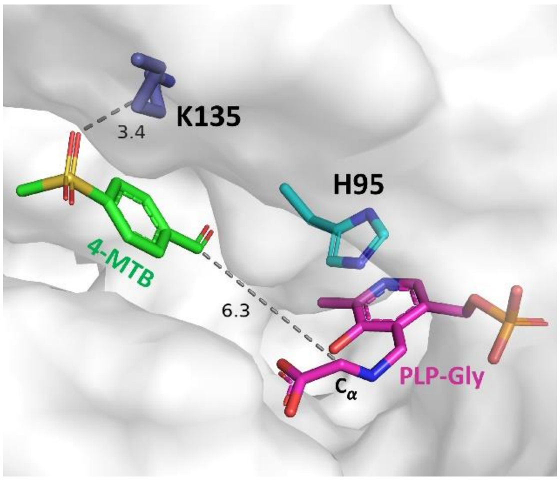

**Figure S7.** Docking models of ternary *Nm*LTA complex in cavity B. PLP-Gly and 4-MTB are shown in magenta and green sticks, respectively. All residues are shown in sticks. Only K135 forms a hydrogen bond interaction with the sulfonic-acid group of the 4-MTB.

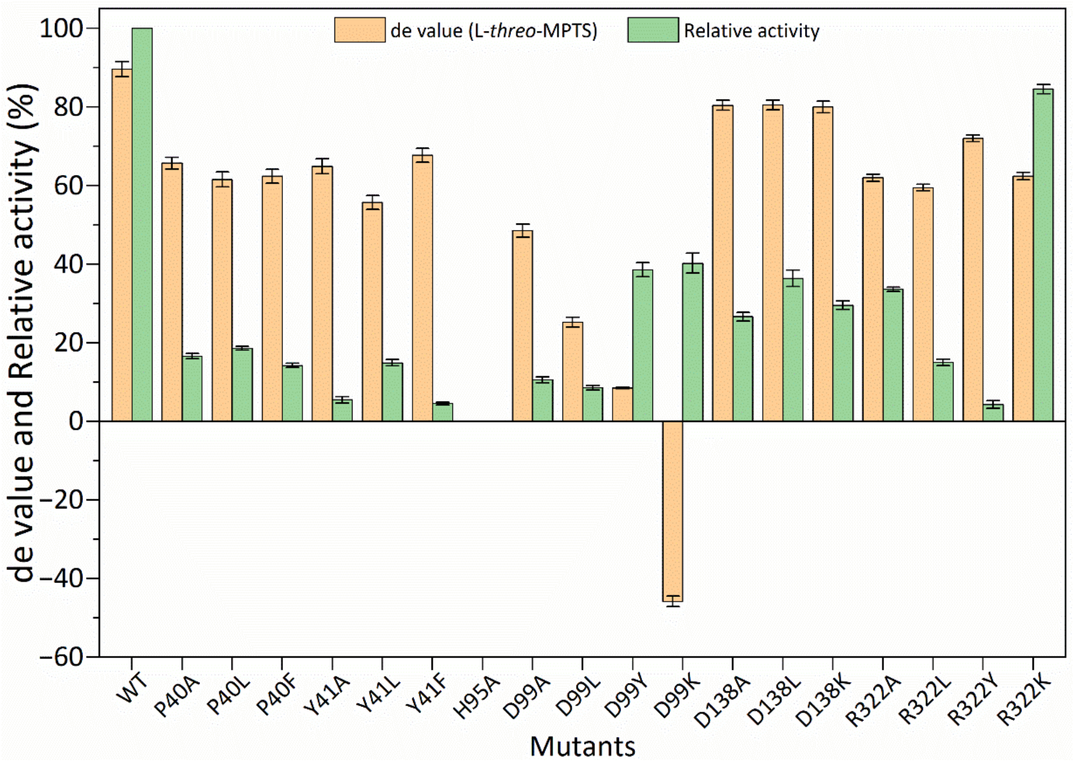

**Figure S8.** The results of P40, Y41, H95, D99, D138 and R322 site-directed mutation. Reaction conditions: 100 mM 4-MTB, 1 M glycine, 50 μM PLP, 10% DMF, and 20 μg purified LTAs in 1mL of 50 mM HEPES buffer (pH 8.0) at 25 °C and 250 rpm within 30 min. All experiments were conducted in triplicate.

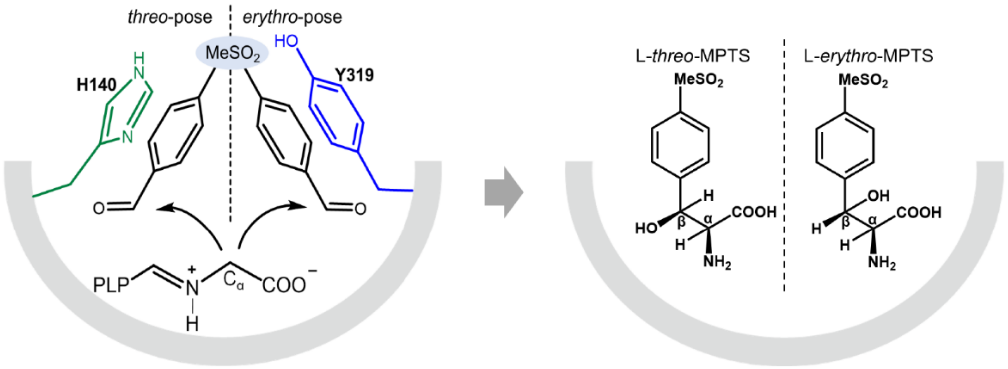

**Figure S9.** "Dual-conformation" regulation mechanism for the diastereoselectivity control of *Nm*LTA. Two distinct binding orientations of 4-MTB lead to the formation of two different HAAs configurations, i.e., L-*threo*-MPTS or L-*erythro*-MPTS.

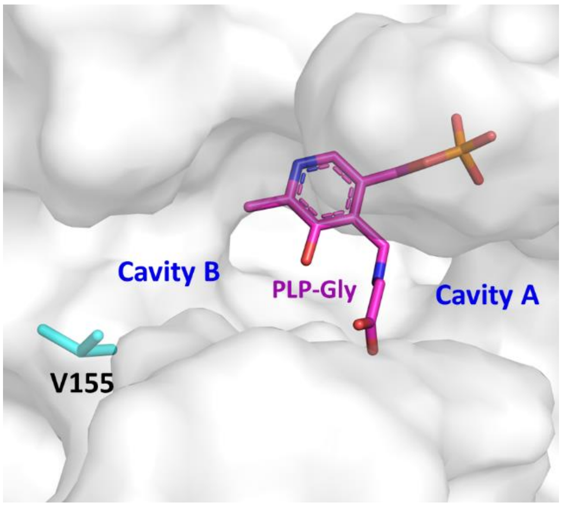

**Figure S10.** Highlight of the V155 residue in cavity B of *Nm*LTA.

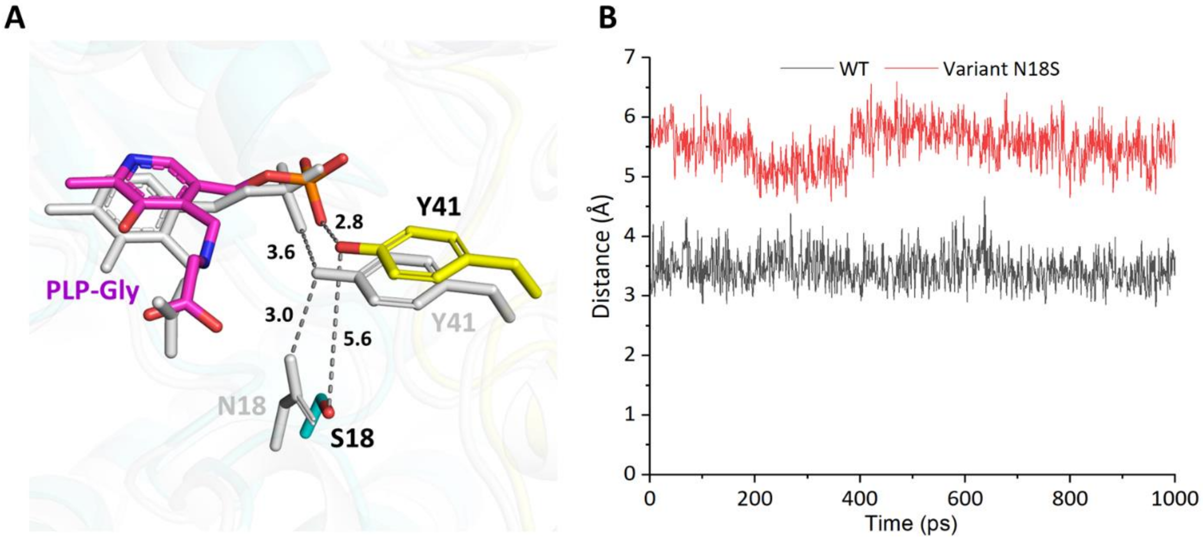

**Figure S11.** Comparison of WT *Nm*LTA with N18S variant. (A) Overlay of *Nm*LTA and N18S variant obtained from constrained- molecular dynamic simulations. Residues are shown in sticks. PLP is shown in stick and colored in magenta (N18S) and gray (WT). (B) The distance plots between N18/S18 and OH-group of Y41 during the initial 1000 ps of molecular dynamics simulations for WT *Nm*LTA and N18S variant.

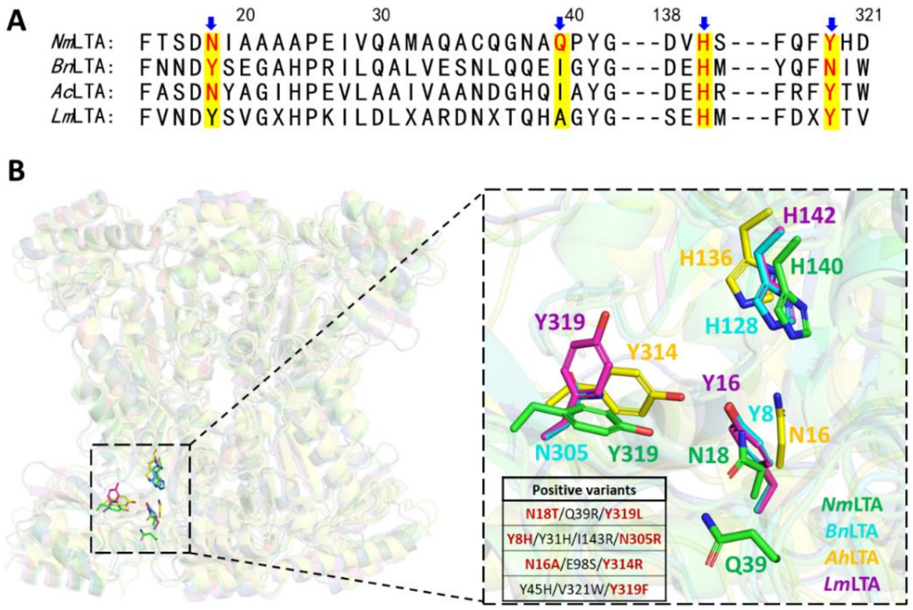

**Figure S12**. Comparison of *Nm*LTA and reported three LTAs mutation sites. (A) Amino acid sequence alignment of four LTAs. The amino acid residues of *Nm*LTA are numbered. (B) Structural comparison of mutation sites of four LTAs from different sources. *Nm*LTA (*Neptunomonas marina*), *Bn*LTA (*Bacillus nealsonii*), *Ah*LTA (*Actinocorallia herbida*), and *Lm*LTA (*Leishmania major*). All residues are shown in sticks. *Nm*LTA, *Bn*LTA, *Ah*LTA, and *Lm*LTA are colored in green, cyan, yellow, and magenta, respectively.

### 1.4 Supporting Tables

**Table S1.** Conversions and diastereoselectivities of 8 candidate LTAs for the synthesis of L-*threo*-MPTS.

| Name | Source | Sequence ID | de value (%) | Conversion (%) |
|---|---|---|---|---|
| *Ml*LTA | *Mameliella* sp. | WP_088714485.1 | 61.8 | 1.0 |
| *Sp*LTA | *Sulfitobacter pontiacus* | HBR36967.1 | 56.3 | 13.1 |
| *Al*LTA | *Actibacterium lipolyticum* | WP_093965798.1 | 51.9 | 0.3 |
| *Nm*LTA | *Neptunomonas marina* | WP_127693843.1 | 89.5 | 38.9 |
| *Rp*LTA | *Rhodopirellula* sp. MGV | WP_094415993.1 | 76.8 | 1.9 |
| *Aj*LTA[17] | *Aeromonas jandaei* DK-39 | O07051.1 | 68.4 | 1.5 |
| *Ec*LTA[18] | *Escherichia coli* | WP_001357131.1 | -6.9 | 0.5 |
| *Pp*LTA[19] | *Pseudomonas putida* | WP_016497628.1 | -41.0 | 0.2 |

Reaction conditions: 100 mM aldehyde, 1 M glycine, 50 µM PLP, 10% DMF, and 20 µg/mL purified LTAs in 1mL of 50 mM HEPES-NaOH buffer (pH 8.0) at 25 °C and 250 rpm within 30 min.

**Table S2.** Kinetic parameters of *Nm*LTA with natural substrates.

| Entry | Substrate | $k_{cat}$ ($s^{-1}$) | $K_m$ (mM) | $k_{cat}/K_m$ ($s^{-1}mM^{-1}$) |
|---|---|---|---|---|
| 1 | L-threonine | 805.7 ± 30.8 | 12.3 ± 0.6 | 65.5 ± 3.1 |
| 2 | L-*allo*-threonine | 1502.4 ± 50.8 | 4.8 ± 0.2 | 317.6 ± 12.5 |
| 3 | D-threonine | n.d. | n.d. | n.d. |
| 4 | D-*allo*-threonine | n.d. | n.d. | n.d. |

The reaction conditions: 0.5 - 50 mM D/L-threonine, 50 μM PLP, 2 mM NADH, 10 U ADH, and 10 μg purified LTAs in 1mL of 50 mM HEPES buffer (pH 8.0) at 25°C. All experiments were conducted in triplicate.

n.d.: not detected.

**Table S3.** Properties of WT *Nm*LTA with different cosolvents.

| Entry | Co-solvent | Co-solvent (v/v) % | Relative activity (%) | 1 h | |
|---|---|---|---|---|---|
| | | | | Conv. (%) | de (%) |
| 1 | Control | - | 100 | 30.5 | 85.6 |
| 2 | DMSO | 5 | 109 | 33.1 | 89.1 |
| 3 | | 10 | 164 | 49.0 | 89.5 |
| 4 | | 15 | 156 | 47.6 | 89.2 |
| 5 | | 20 | 145 | 44.2 | 88.5 |
| 6 | | 25 | 137 | 41.8 | 88.3 |
| 7 | DMF | 5 | 115 | 33.4 | 89.0 |
| 8 | | 10 | 168 | 51.2 | 89.8 |
| 9 | | 15 | 154 | 47.0 | 89.3 |
| 10 | | 20 | 149 | 45.4 | 89.1 |
| 11 | | 25 | 142 | 43.3 | 88.9 |

Reaction conditions: 100 mM 4-MTB, 1 M glycine, 50 mM PLP, and 1.5 U LTAs in 1mL of 50 mM HEPES buffer (pH 8.0) at 25 °C and 250 rpm for 60 min.

**Table S4.** X-Ray crystallographic data collection and refinement statistics.

| | WT *Nm*LTA（PDB ID: 7YVR） |
|---|---|
| **Data collection** | |
| Wavelength | 0.97853 |
| Resolution (Å) | 19.820 - 2.798 (2.898 - 2.798) |
| Space group | C 1 2 1 |
| Cell dimensions | |
| a, b) c (Å) | 264.4, 59.9, 95.1 |
| α, β, γ (°) | 90.0, 106.1, 90.0 |
| Mean I/sigma(I) | 7.65 (2.43) |
| Completeness (%) | 98.97 (99.09) |
| Redundancy | 4.5 (4.4) |
| Wilson B Factor (Å$^2$) | 35.44 |
| R-meas | 0.1602 (0.5213) |
| R-merge | 0.1822 (0.5926) |
| CC1/2 | 0.984 (0.819) |
| **Refinement** | |
| No. reflections | 35429 |
| $R_{work}$ | 0.2070 (0.2828) |
| $R_{free}$ | 0.2391 (0.3451) |
| Number of non-H atoms | 10526 |
| Protein | 10339 |
| Ligand | 18 |
| Water | 169 |
| Average B-factor (Å$^2$) | 34.85 |
| Protein | 34.85 |
| Ligand | 20 |
| Water | 34.2 |
| **Ramachandran plot** | |
| Favored (%) | 97.41 |
| Allowed (%) | 2.59 |
| **R.M.S deviations** | |
| Bond lengths (Å) | 0.008 |
| Bond angles (°) | 1.11 |

**Table S5.** Primers used in this study.

| Name | Primer sequence (5' to 3') [a] |
|---|---|
| P40A-F | AATGCGCAAGCGTACGGCGGAGACGCGCTG |
| P40A-R | AATGCGCAACGCTACGGCGGAGACGCGCTG |
| P40L-F | AATGCGCAACTGTACGGCGGAGACGCGCTG |
| P40L-R | AATGCGCAACAGTACGGCGGAGACGCGCTG |
| P40F-F | AATGCGCAATTCTACGGCGGAGACGCGCTG |
| P40F-R | AATGCGCAAGAATACGGCGGAGACGCGCTG |
| Y41A-F | GCGCAACCGGCGGGCGGAGACGCGCTGACT |
| Y41A-R | GTCTCCGCCCGCCGGTTGCGCATTGCCCTG |
| Y41L-F | GCGCAACCGCTGGGCGGAGACGCGCTGACT |
| Y41L-R | GTCTCCGCCCAGCGGTTGCGCATTGCCCTG |
| Y41F-F | GCGCAACCGTTCGGCGGAGACGCGCTGACT |
| Y41F-R | GTCTCCGCCGAACGGTTGCGCATTGCCCTG |
| H95A-F | CAGGAGAGCGCGATTAACAACGATGAGTGC |
| H95A-R | GTTGTTAATCGCGCTCTCCTGGTGGCACAA |
| D99A-F | ATTAACAACGCGGAGTGCGGTGCGCCGGAA |
| D99A-R | ACCGCACTCCGCGTTGTTAATATGGCTCTC |
| D99L-F | ATTAACAACCTGGAGTGCGGTGCGCCGGAA |
| D99L-R | ACCGCACTCCAGGTTGTTAATATGGCTCTC |
| D99Y-F | ATTAACAACTATGAGTGCGGTGCGCCGGAA |
| D99Y-R | ACCGCACTCATAGTTGTTAATATGGCTCTC |
| D99K-F | ATTAACAACAAGGAGTGCGGTGCGCCGGAA |
| D99K-R | ACCGCACTCCTTGTTGTTAATATGGCTCTC |
| D138A-F | AAACGCGGCGCGGTTCACAGCGTCGAGCCG |
| D138A-R | GCTGTGAACCGCGCCGCGTTTATTGCGCGC |
| D138L-F | AAACGCGGCCTGGTTCACAGCGTCGAGCCG |
| D138L-R | GCTGTGAACCAGGCCGCGTTTATTGCGCGC |
| D138K-F | AAACGCGGCAAGGTTCACAGCGTCGAGCCG |
| D138K-R | GCTGTGAACCTTGCCGCGTTTATTGCGCGC |
| R322A-F | TATCACGATGCGTGGGGCGACGGCATTGTT |
| R322A-R | GTCGCCCCACGCATCGTGATAAAACTGAAA |
| R322L-F | TATCACGATCTGTGGGGCGACGGCATTGTT |
| R322L-R | GTCGCCCCACAGATCGTGATAAAACTGAAA |
| R322Y-F | TATCACGATTATTGGGGCGACGGCATTGTT |
| R322Y-R | GTCGCCCCAATAATCGTGATAAAACTGAAA |
| R322K-F | TATCACGATAAGTGGGGCGACGGCATTGTT |

| | |
|---|---|
| R322K-R | GTCGCCCCA<u>CTT</u>ATCGTGATAAAACTGAA |
| S16-F | AGCTTCACC<u>NNK</u>GACAACATCGCGGCTGCG |
| S16-R | GATGTTGTC<u>MNN</u>GGTGAAGCTCACTGTGTC |
| N18-F | ACCTCCGAC<u>NNK</u>ATCGCGGCTGCGGCTCCG |
| N18-R | AGCCGCGAT<u>MNN</u>GTCGGAGGTGAAGCTCAC |
| Q39-F | GGCAATGCG<u>NNK</u>CCGTACGGCGGAGACGCG |
| Q39-R | GCCGTACGG<u>MNN</u>CGCATTGCCCTGACACGC |
| H140-F | GGCGACGTT<u>NNK</u>AGCGTCGAGCCGACCACC |
| H140-R | CTCGACGCT<u>MNN</u>AACGTCGCCGCGTTTATT |
| Y319-F | TTTCAGTTT<u>NNK</u>CACGATCGTTGGGGCGAC |
| Y319-R | ACGATCGTG<u>MNN</u>AAACTGAAAACCCTGTTG |
| S16A-F | AGCTTCACC<u>GCG</u>GACAACATCGCGGCTGCG |
| S16A-R | GATGTTGTC<u>CGC</u>GGTGAAGCTCACTGTGTC |
| S16G-F | AGCTTCACC<u>GGC</u>GACAACATCGCGGCTGCG |
| S16G-R | GATGTTGTC<u>GCC</u>GGTGAAGCTCACTGTGTC |
| N18S-F | ACCTCCGAC<u>AGC</u>ATCGCGGCTGCGGCTCCG |
| N18S-R | AGCCGCGAT<u>GCT</u>GTCGGAGGTGAAGCTCAC |
| N18T-F | ACCTCCGAC<u>ACG</u>ATCGCGGCTGCGGCTCCG |
| N18T-R | AGCCGCGAT<u>CGT</u>GTCGGAGGTGAAGCTCAC |
| Q39R-F | GGCAATGCG<u>CGT</u>CCGTACGGCGGAGACGCG |
| Q39R-R | GCCGTACGG<u>ACG</u>CGCATTGCCCTGACACGC |
| Y319L-F | TTTCAGTTT<u>CTG</u>CACGATCGTTGGGGCGAC |
| Y319L-R | ACGATCGTG<u>CAG</u>AAACTGAAAACCCTGTTG |
| Y319D-F | TTTCAGTTT<u>GAC</u>CACGATCGTTGGGGCGAC |
| Y319D-R | ACGATCGTG<u>GTC</u>AAACTGAAAACCCTGTTG |

[a] N is A, G, C, or T; K is G or T; M is A or C.

**Table S6.** Conversion and de value of the WT *Nm*LTA positive mutants produced.

| Entry | Enzyme | Specific Activity （U/mg） | de (%) | Conv. (%) |
|---|---|---|---|---|
| 1 | WT *Nm*LTA | 64.8 | 89.6 | 81.3 |
| 2 | S16A | 29.3 | 94.3 | 81.4 |
| 3 | S16G | 16.3 | 94.1 | 78.3 |
| 4 | N18T | 48.0 | 91.1 | 80.6 |
| 5 | N18S | 43.2 | 92.5 | 83.8 |
| 6 | Q39A | 60.5 | 92.3 | 85.7 |
| 7 | Q39R | 68.3 | 92.6 | 80.9 |
| 8 | Q39K | 72.8 | 91.8 | 83.2 |
| 9 | Q39H | 65.4 | 91.6 | 77.5 |
| 10 | Q39L | 60.7 | 92.2 | 80.3 |
| 11 | Y319L | 55.9 | 93.7 | 84.5 |
| 12 | Y319S | 66.2 | 92.7 | 83.6 |
| 13 | Y319D | 17.8 | 94.0 | 80.9 |
| 14 | S16A/Y319L | 1.3 | 95.8 | 81.4 |
| 15 | S16A/Q39R | 10.5 | 95.1 | 80.6 |
| 16 | Q39R/Y319D | 19.2 | 95.8 | 79.6 |
| 17 | S16G/Y319L | 32.3 | 96.2 | 80.3 |
| 18 | N18S/Y319L | 64.4 | 96.8 | 81.0 |
| 19 | Q39R/Y319L | 133.0 | 97.6 | 81.7 |
| 20 | N18S/Q39R/Y319D | 47.2 | 98.5 | 77.3 |
| 21 | N18S/Q39R/Y319L | 95.7 | 99.3 | 82.6 |

Reaction conditions: 100 mM 4-MTB, 1 M glycine, 50 μM PLP, 10% DMF, and 1.5 U LTAs in 1mL of 50 mM HEPES buffer (pH 8.0) at 25 °C and 250 rpm for 120 min.

**Table S7**. Kinetic parameters of the WT *Nm*LTA and its mutants.

| Entry | Enzyme | Products | $k_{cat}$ ($s^{-1}$) | $K_m$ (mM) | $k_{cat}/K_m$ ($s^{-1}$ $mM^{-1}$) | SP* |
|---|---|---|---|---|---|---|
| 1 | WT *Nm*LTA | L-*threo*-MPTS | 190.4 ± 5.6 | 4.5 ± 0.1 | 42.4 ± 0.3 | 27.8 |
| | | L-*erythro*-MPTS | 12.0 ± 0.5 | 7.9 ± 0.2 | 1.5 ± 0.0 | |
| 2 | N18S | L-*threo*-MPTS | 155.3 ± 3.2 | 6.5 ± 0.2 | 23.9 ± 0.2 | 57.8 |
| | | L-*erythro*-MPTS | 2.2 ± 0.1 | 5.4 ± 0.1 | 0.4 ± 0.0 | |
| 3 | Q39R | L-*threo*-MPTS | 207.6 ± 5.9 | 4.8 ± 0.3 | 43.2 ± 0.4 | 72.6 |
| | | L-*erythro*-MPTS | 1.7 ± 0.1 | 2.8 ± 0.1 | 0.6 ± 0.0 | |
| 4 | Y319L | L-*threo*-MPTS | 146.2 ± 3.4 | 5.0 ± 0.2 | 29.2 ± 0.3 | 43.3 |
| | | L-*erythro*-MPTS | 3.5 ± 0.1 | 5.2 ± 0.1 | 0.7 ± 0.0 | |
| 5 | Q39R/Y319L | L-*threo*-MPTS | 400.3 ± 10.6 | 5.0 ± 0.2 | 80.2 ± 0.6 | 160.1 |
| | | L-*erythro*-MPTS | 3.6 ± 0.1 | 7.2 ± 0.3 | 0.5 ± 0.01 | |
| 6 | N18S/Q39R/Y319L | L-*threo*-MPTS | 315.4 ± 9.5 | 4.0 ± 0.6 | 78.9 ± 0.5 | 297.5 |
| | | L-*erythro*-MPTS | 2.2 ± 0.02 | 8.3 ± 0.4 | 0.3 ± 0.02 | |

Reaction conditions: 0.1-100 mM 4-MTB, 1-1000 mM glycine, 50 μM PLP, 10% DMF, and 10 μg purified LTAs in 1mL of HEPES buffer (50 mM, pH 8.0) at 25 °C and 250 rpm within 10 min. All experiments were conducted in triplicate.

*SP is the ratio of $(k_{cat}/K_m)_{threo}$ to $(k_{cat}/K_m)_{erythro}$.

### 1.5 Results of HPLC and MS

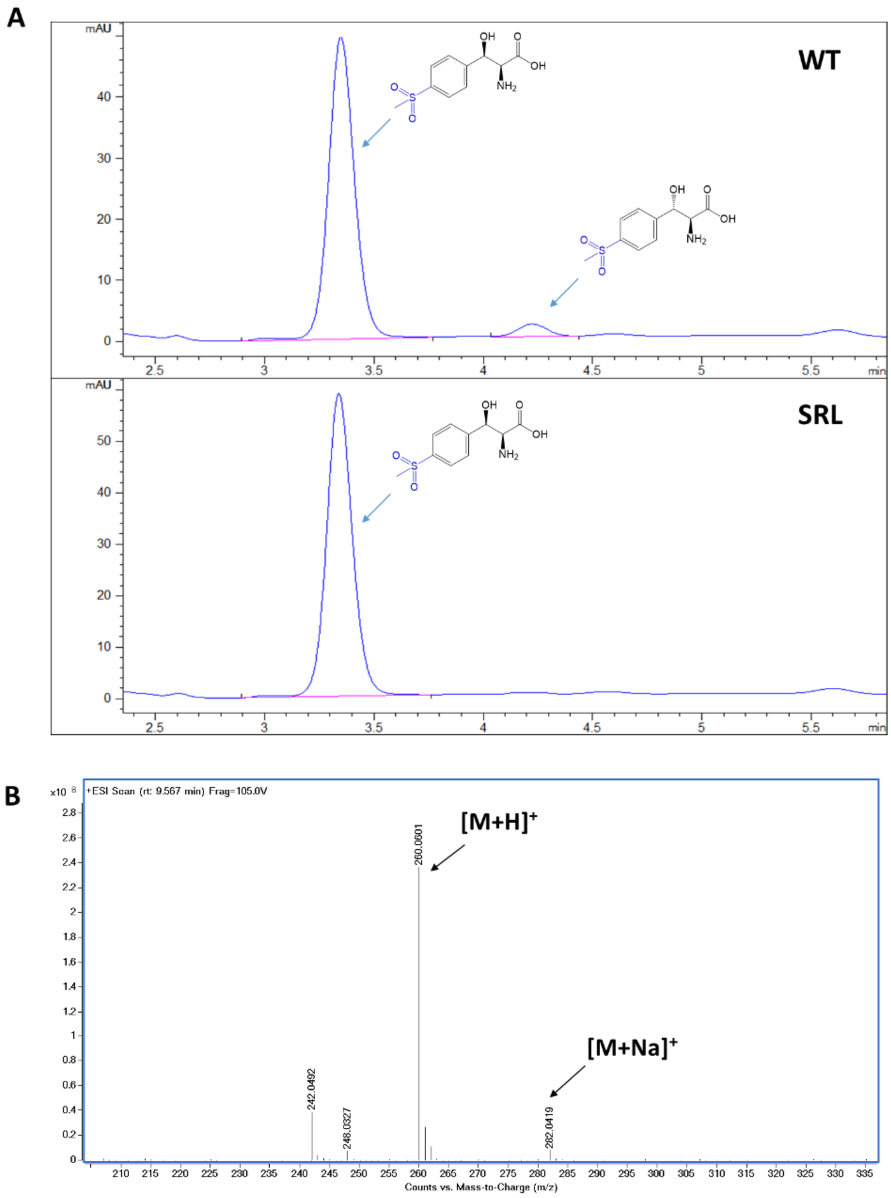

**Figure S13.** HPLC and MS analysis of L-4-methylsulfonylphenylserine catalyzed by WT *Nm*LTA and the SRL variant. (A) HPLC analysis of L-*threo*-4-methylsulfonylphenylserine and L-4-*erythro*-methylsulfonylphenylserine. The retention time of L-*threo*-4-methylsulfonylphenylserine and L-4-*erythro*-methylsulfonylphenylserine were 3.352 min and 4.216 min, respectively. (B) An ESI-MS spectrum of the product with $[M+H]^+$ at m/z 260.0601 and $[M+Na]^+$ at m/z 282.0419.

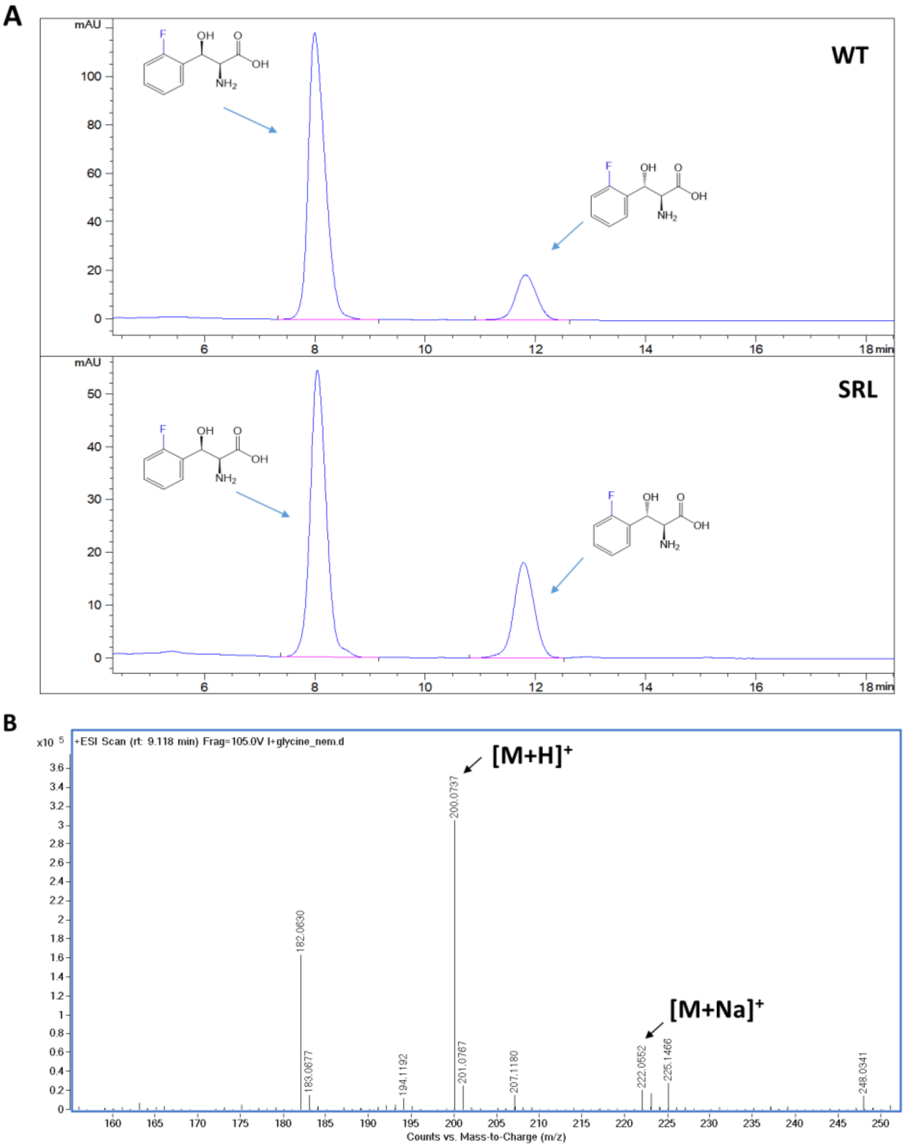

**Figure S14.** HPLC and MS analysis of L-2-F-phenylserine catalyzed by WT *Nm*LTA and the SRL variant. (A) HPLC analysis of L-*threo*-2-F-phenylserine and L-*erythro*-2-F-phenylserine. The retention time of L-*threo*-2-F-phenylserine and L-*erythro*-2-F-phenylserine were 8.005 min and 11.820 min, respectively. (B) An ESI-MS spectrum of the product with $[M+H]^+$ at m/z 200.0737 and $[M+Na]^+$ at m/z 222.0552.

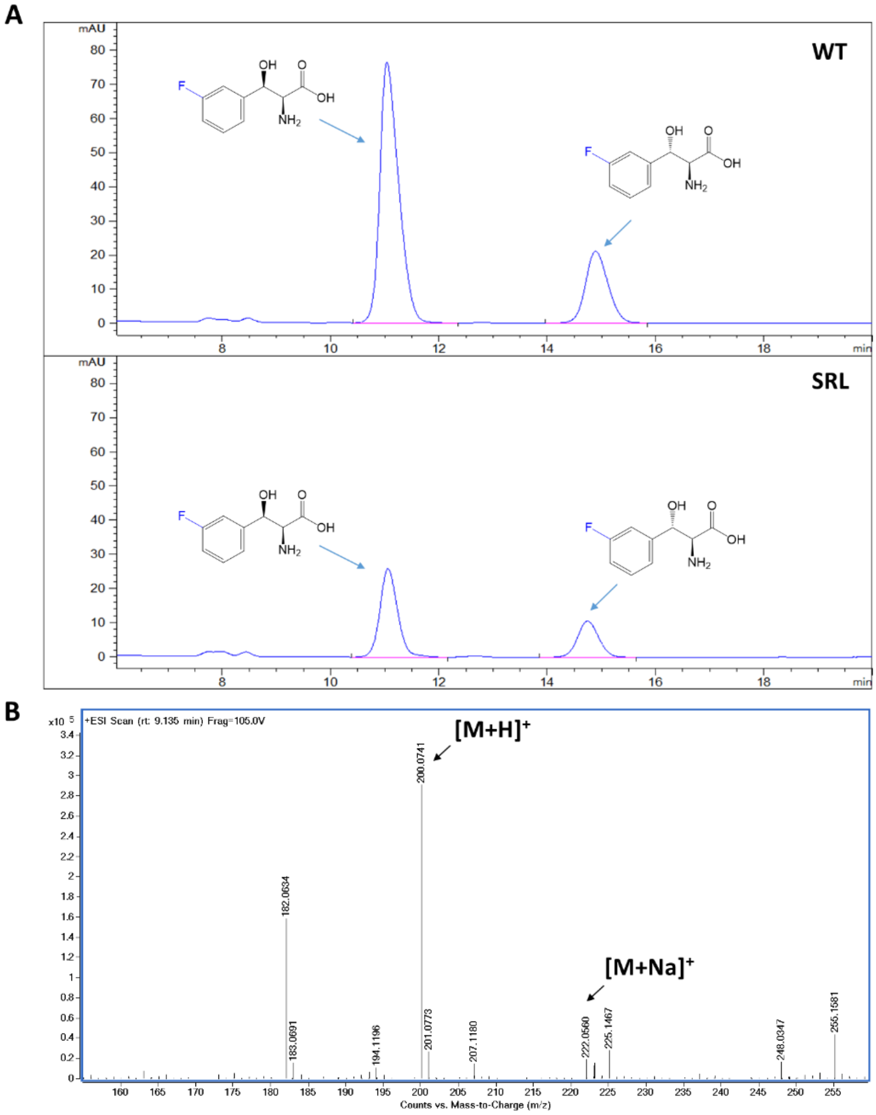

**Figure S15.** HPLC and MS analysis of L-3-F-phenylserine catalyzed by WT *Nm*LTA and the SRL variant. (A) HPLC analysis of L-*threo*-3-F-phenylserine and L-*erythro*-3-F-phenylserine. The retention time of L-*threo*-3-F-phenylserine and L-*erythro*-3-F-phenylserine were 10.010 min and 14.825 min, respectively. (B) An ESI-MS spectrum of the product with $[M+H]^+$ at m/z 200.0741 and $[M+Na]^+$ at m/z 222.0560.

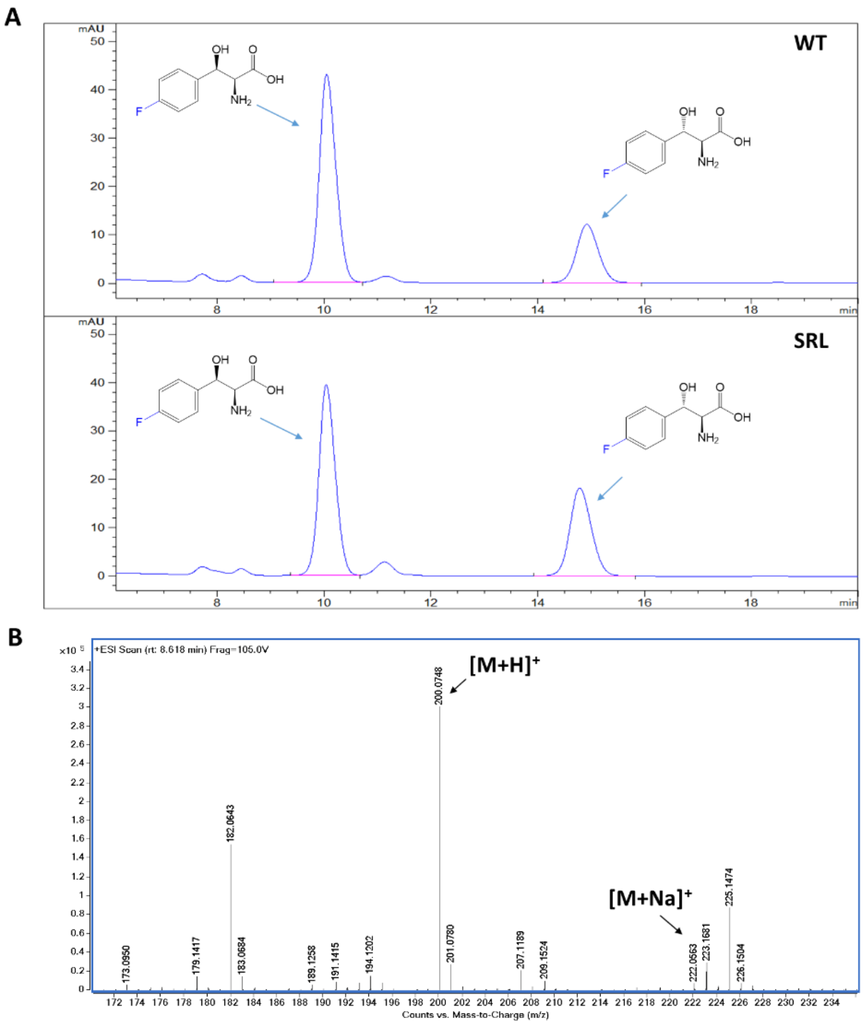

**Figure S16.** HPLC and MS analysis of L-4-F-phenylserine catalyzed by WT *Nm*LTA and the SRL variant. (A) HPLC analysis of L-*threo*-4-F-phenylserine and L-*erythro*-4-F-phenylserine. The retention time of L-*threo*-4-F-phenylserine and L-*erythro*-4-F-phenylserine were 10.050 min and 14.850 min, respectively. (B) An ESI-MS spectrum of the product with $[M+H]^+$ at m/z 200.0748 and $[M+Na]^+$ at m/z 222.0563.

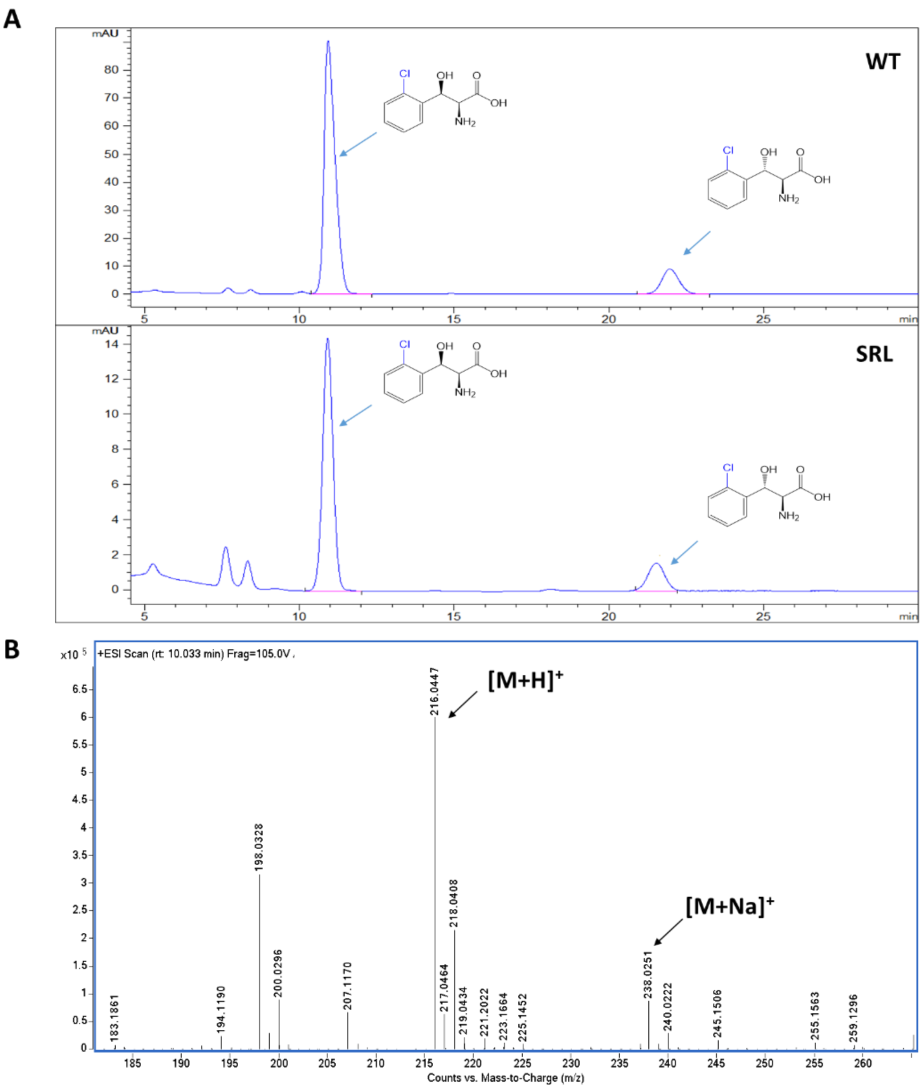

**Figure S17.** HPLC and MS analysis of L-2-Cl-phenylserine catalyzed by WT *Nm*LTA and the SRL variant. (A) HPLC analysis of L-*threo*-2-Cl-phenylserine and L-*erythro*-2-Cl-phenylserine. The retention time of L-*threo*-2-Cl-phenylserine and L-*erythro*-2-Cl-phenylserine were 10.926 min and 21.958 min, respectively. (B) An ESI-MS spectrum of the product with $[M+H]^+$ at m/z 216.0447 and $[M+Na]^+$ at m/z 238.0251.

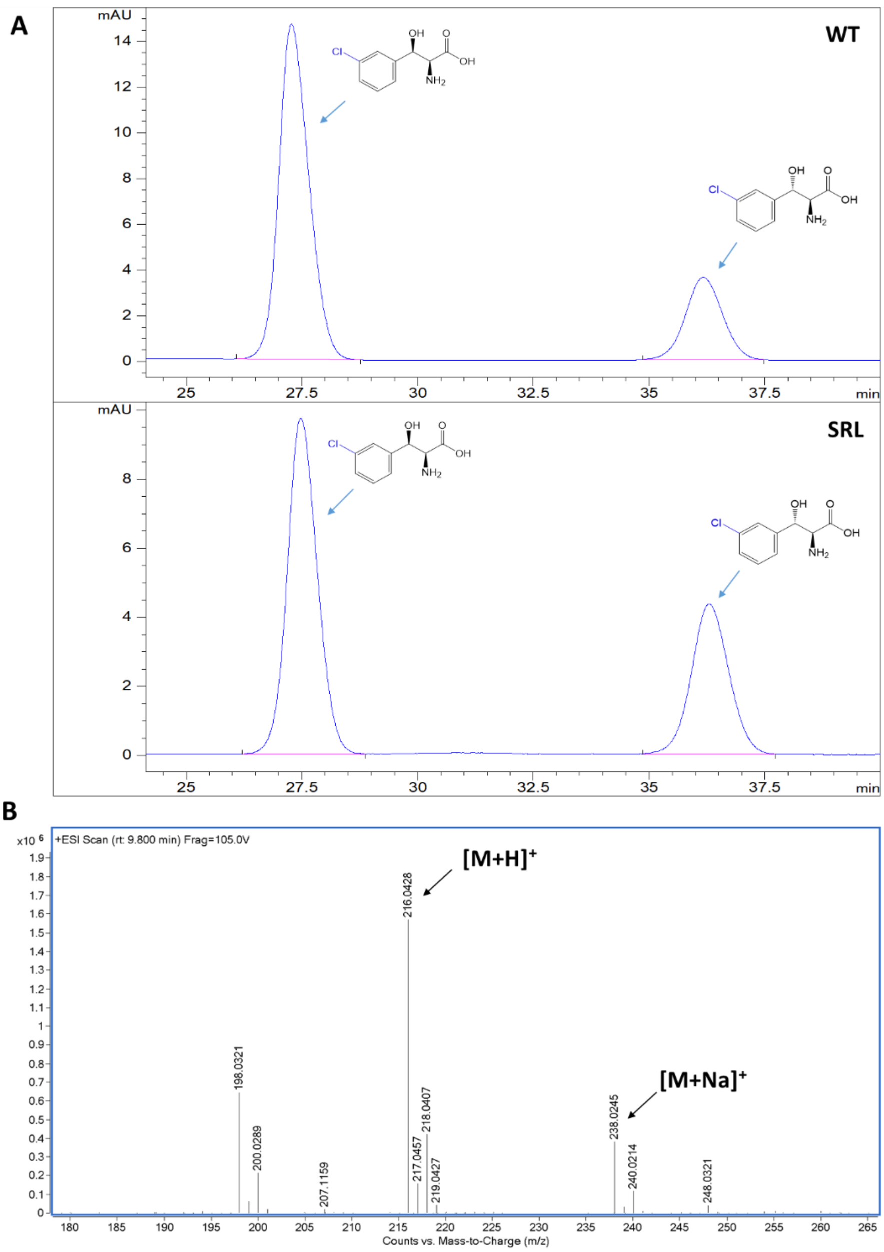

**Figure S18.** HPLC and MS analysis of L-3-Cl-phenylserine catalyzed by WT *Nm*LTA and the SRL variant. (A) HPLC analysis of L-*threo*-3-Cl-phenylserine and L-*erythro*-3-Cl-phenylserine. The retention time of L-*threo*-3-Cl-phenylserine and L-*erythro*-3-Cl-phenylserine were 27.545 min and 36.842 min, respectively. (B) An ESI-MS spectrum of the product with $[M+H]^+$ at m/z 216.0428 and $[M+Na]^+$ at m/z 238.0245.

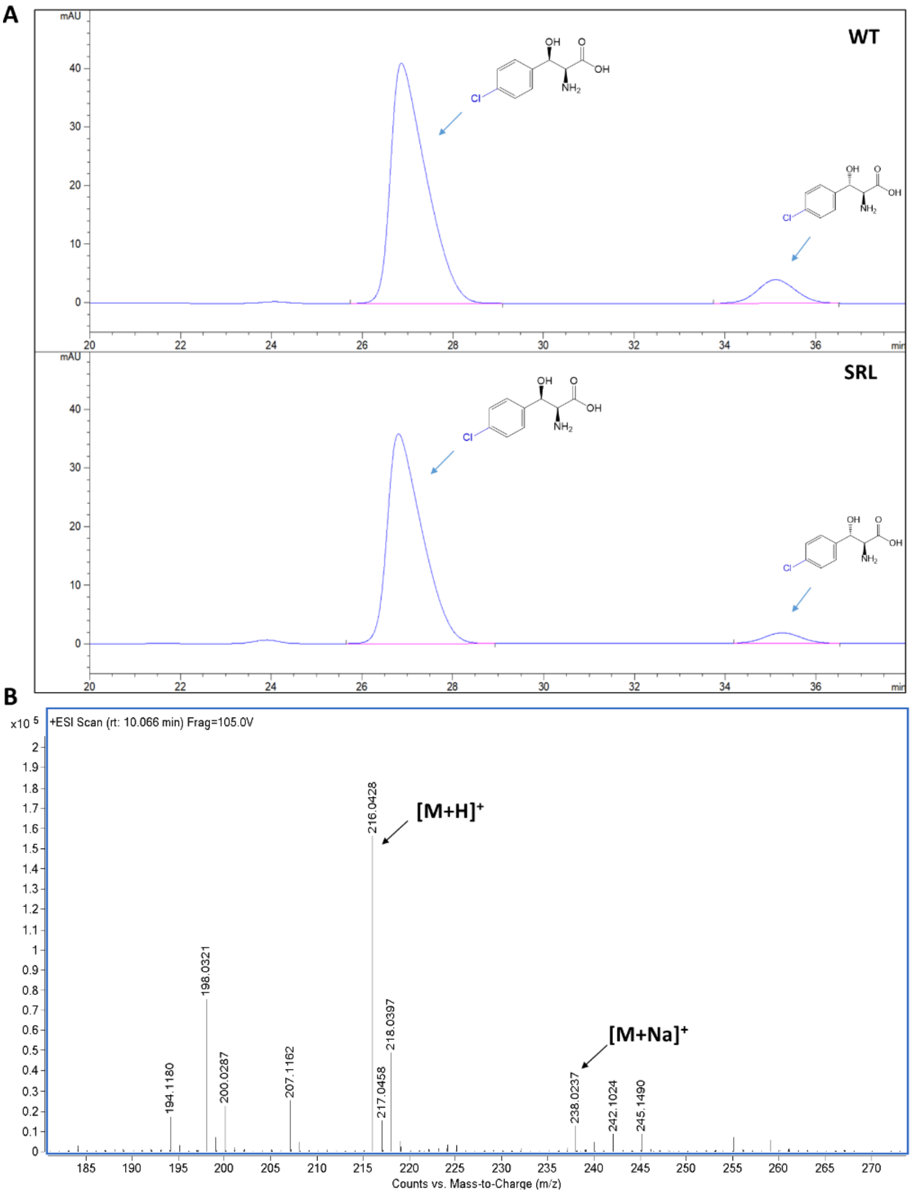

**Figure S19.** HPLC and MS analysis of L-4-Cl-phenylserine catalyzed by WT *Nm*LTA and the SRL variant. (A) HPLC analysis of L-*threo*-4-Cl-phenylserine and L-*erythro*-4-Cl-phenylserine. The retention time of L-*threo*-4-Cl-phenylserine and L-*erythro*-4-Cl-phenylserine were 26.864 min and 35.123 min, respectively. (B) An ESI-MS spectrum of the product with $[M+H]^+$ at m/z 216.0428 and $[M+Na]^+$ at m/z 238.0237.

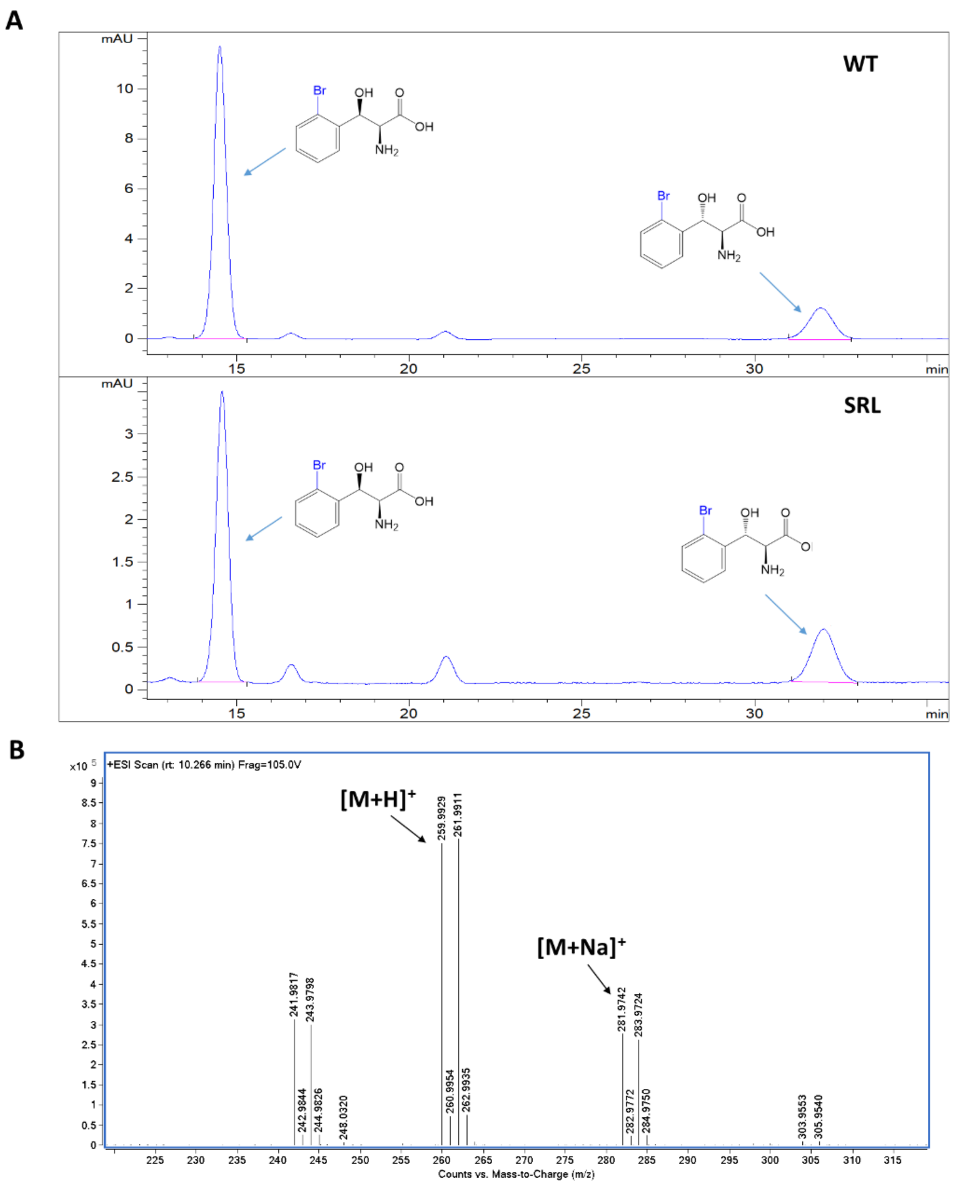

**Figure S20.** HPLC and MS analysis of L-2-Br-phenylserine catalyzed by WT *Nm*LTA and the SRL variant. (A) HPLC analysis of L-*threo*-2-Br-phenylserine and L-*erythro*-2-Br-phenylserine. The retention time of L-*threo*-2-Br-phenylserine and L-*erythro*-2-Br-phenylserine were 14.515 min and 31.922 min, respectively. (B) An ESI-MS spectrum of the product with $[M+H]^+$ at m/z 259.9929 and $[M+Na]^+$ at m/z 281.9742.

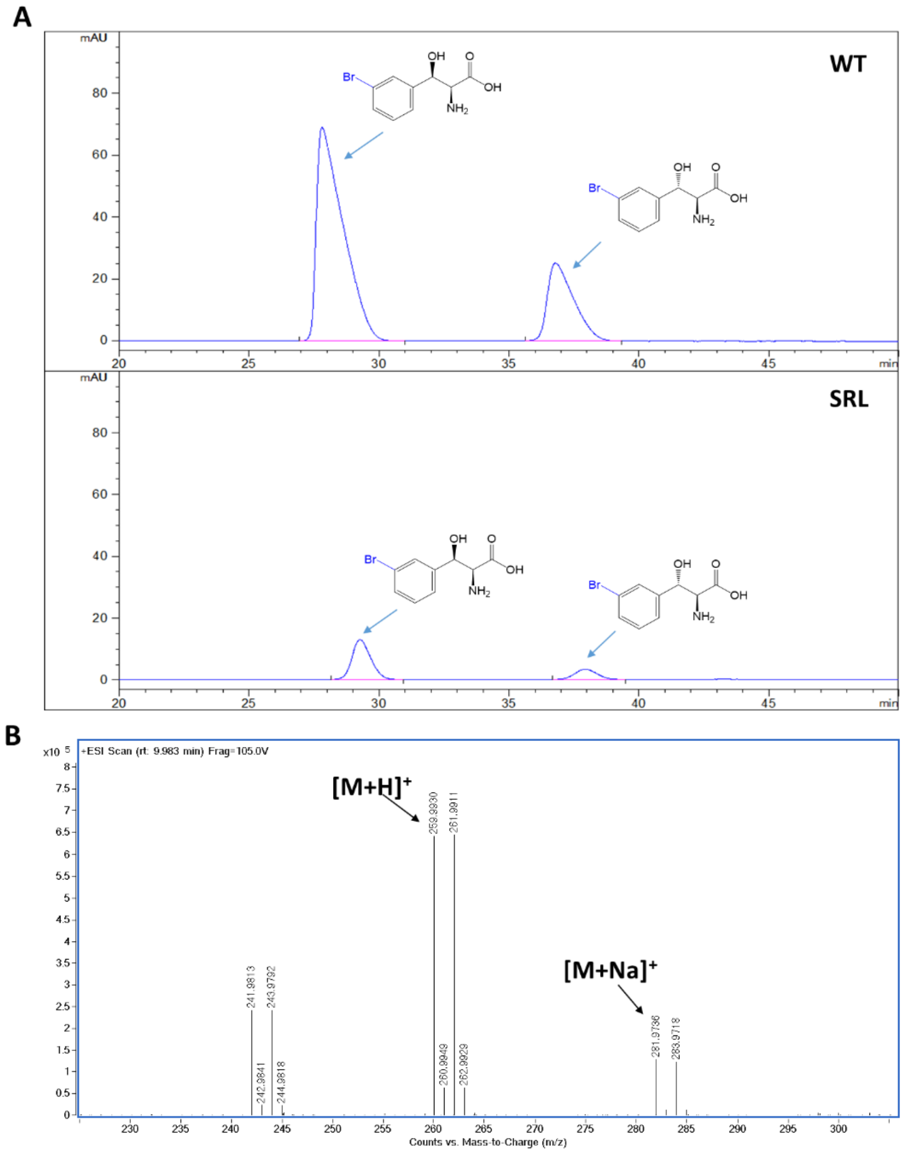

**Figure S21.** HPLC and MS analysis of L-3-Br-phenylserine catalyzed by WT *Nm*LTA and the SRL variant. (A) HPLC analysis of L-*threo*-3-Br-phenylserine and L-*erythro*-3-Br-phenylserine. The retention time of L-*threo*-3-Br-phenylserine and L-*erythro*-3-Br-phenylserine were 27.815 min and 36.794 min, respectively. (B) An ESI-MS spectrum of the product with $[M+H]^+$ at m/z 259.9930 and $[M+Na]^+$ at m/z 281.9736.

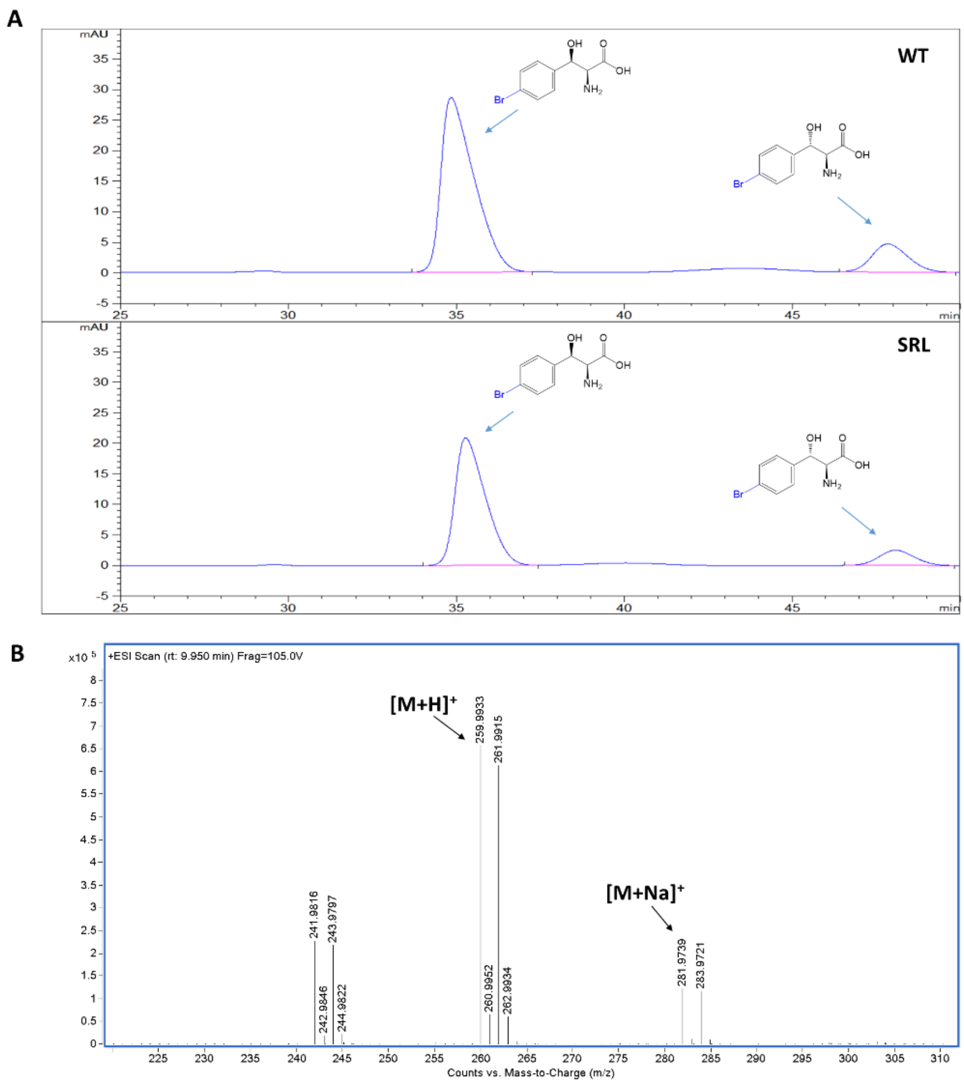

**Figure S22.** HPLC and MS analysis of L-4-Br-phenylserine catalyzed by WT *Nm*LTA and the SRL variant. (A) HPLC analysis of L-*threo*-4-Br-phenylserine and L-*erythro*-4-Br-phenylserine. The retention time of L-*threo*-4-Br-phenylserine and L-*erythro*-4-Br-phenylserine were 34.842 min and 47.850 min, respectively. (B) An ESI-MS spectrum of the product with $[M+H]^+$ at m/z 259.9933 and $[M+Na]^+$ at m/z 281.9739.

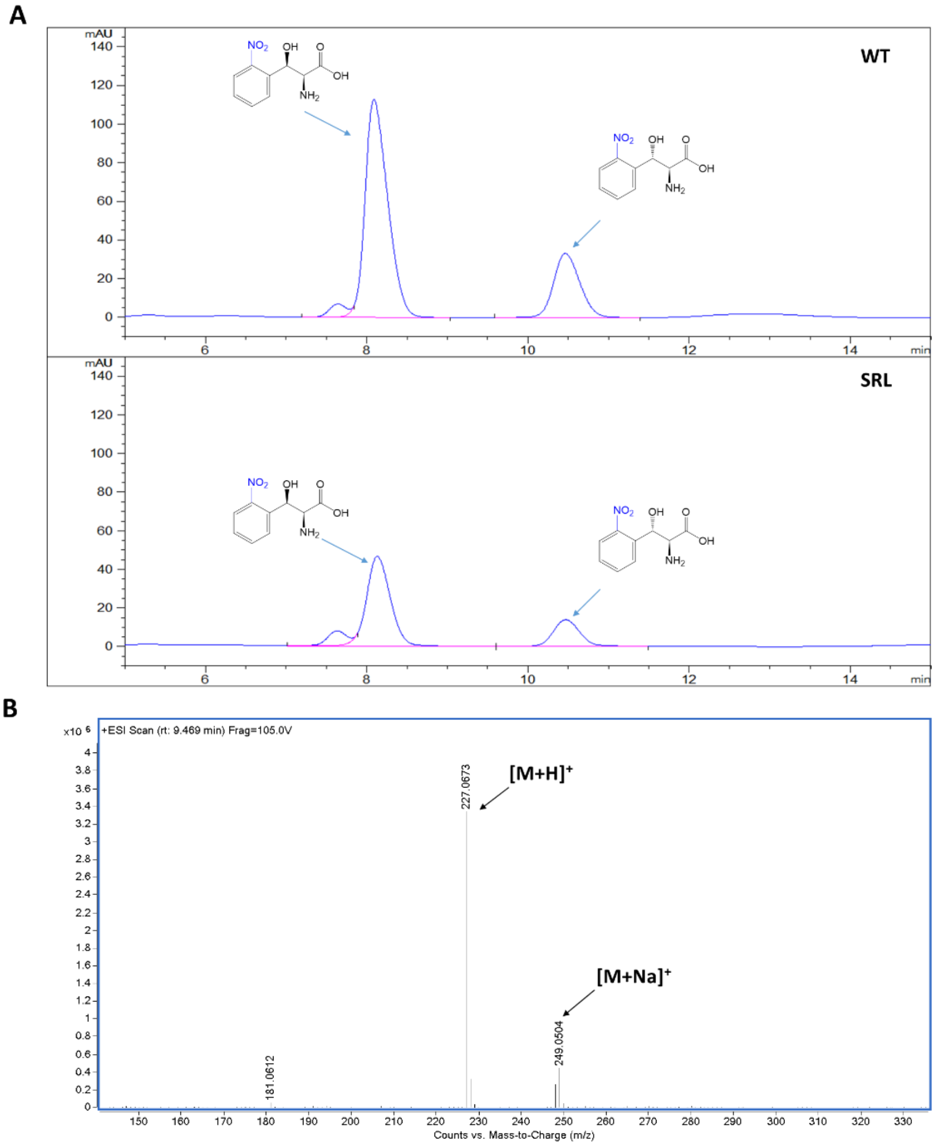

**Figure S23.** HPLC and MS analysis of L-2-$NO_2$-phenylserine catalyzed by WT *Nm*LTA and the SRL variant. (A) HPLC analysis of L-*threo*-2-$NO_2$-phenylserine and L-*erythro*-2-$NO_2$-phenylserine. The retention time of L-*threo*-2-$NO_2$-phenylserine and L-*erythro*-2-$NO_2$-phenylserine were 8.091 min and 10.446 min, respectively. (B) An ESI-MS spectrum of the product with $[M+H]^+$ at m/z 227.0673 and $[M+Na]^+$ at m/z 249.0504.

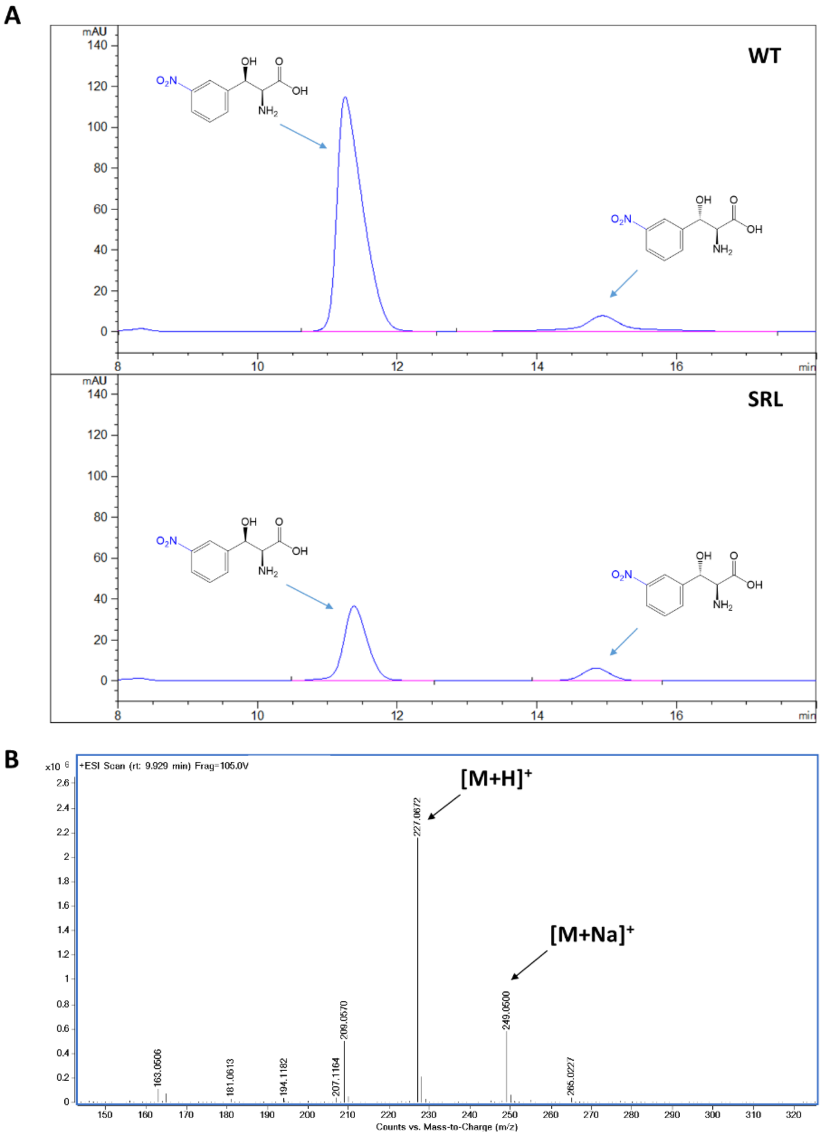

**Figure S24.** HPLC and MS analysis of L-3-$NO_2$-phenylserine catalyzed by WT *Nm*LTA and the SRL variant. (A) HPLC analysis of L-*threo*-3-$NO_2$-phenylserine and L-*erythro*-3-$NO_2$-phenylserine. The retention time of L-*threo*-3-$NO_2$-phenylserine and L-*erythro*-3-$NO_2$-phenylserine were 11.253 min and 14.939 min, respectively. (B) An ESI-MS spectrum of the product with $[M+H]^+$ at m/z 227.0672 and $[M+Na]^+$ at m/z 249.0500.

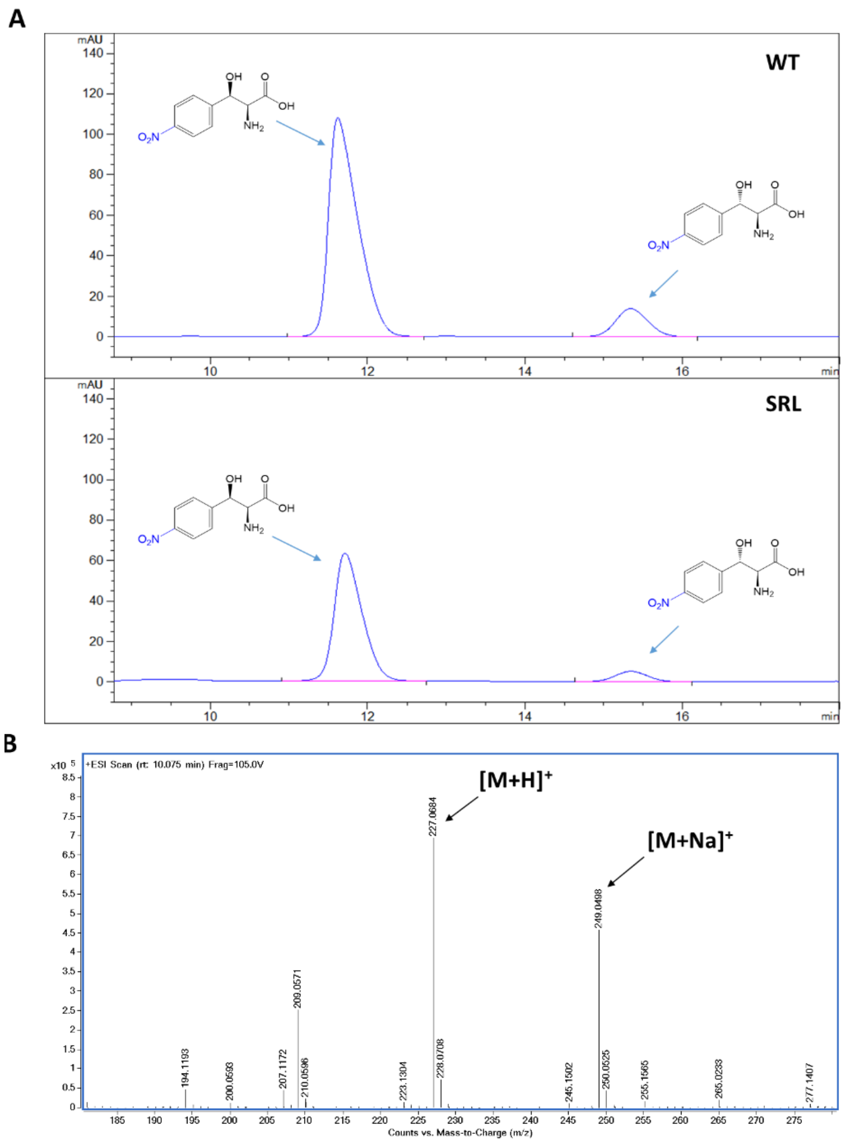

**Figure S25.** HPLC and MS analysis of L-4-$NO_2$-phenylserine catalyzed by WT *Nm*LTA and the SRL variant. (A) HPLC analysis of L-*threo*-4-$NO_2$-phenylserine and L-*erythro*-4-$NO_2$-phenylserine. The retention time of L-*threo*-4-$NO_2$-phenylserine and L-*erythro*-4-$NO_2$-phenylserine were 11.622 min and 15.344 min, respectively. (B) An ESI-MS spectrum of the product with $[M+H]^+$ at m/z 227.0684 and $[M+Na]^+$ at m/z 249.0498.

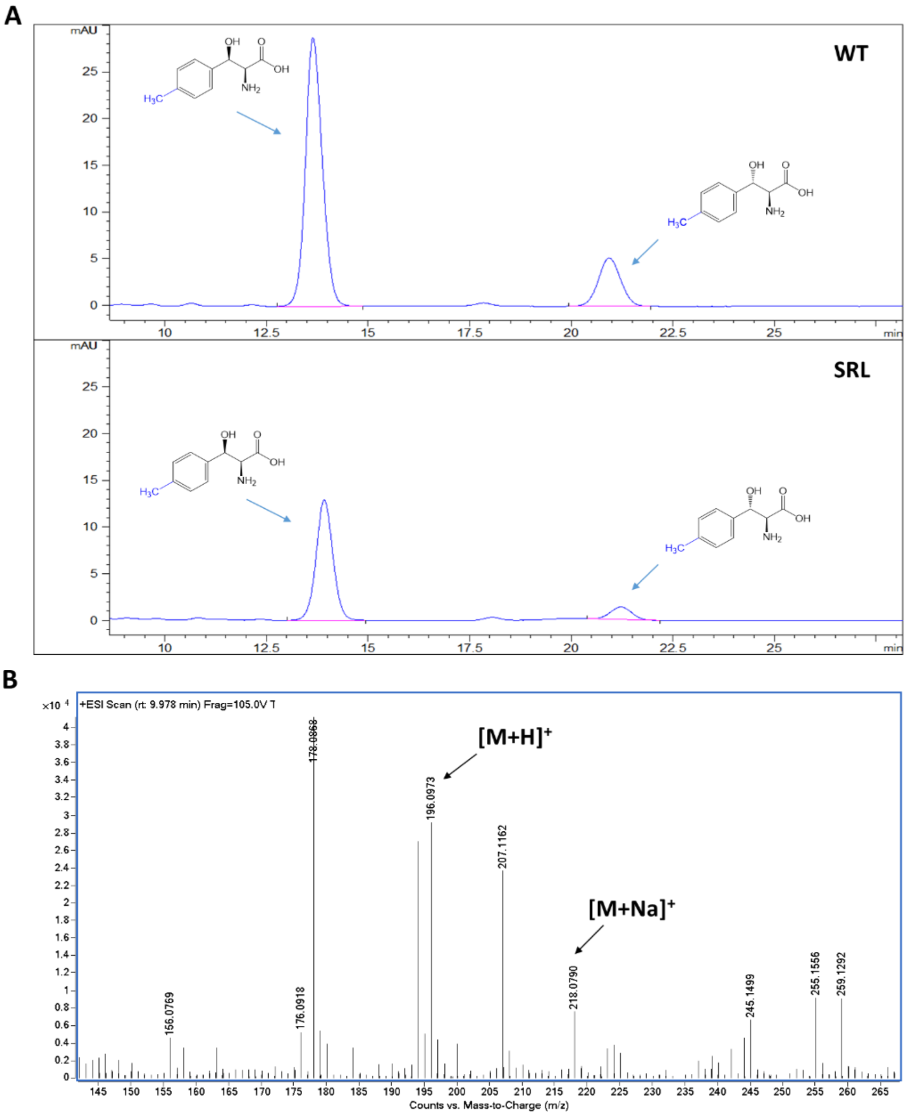

**Figure S26.** HPLC and MS analysis of L-4-$CH_3$-phenylserine catalyzed by WT *Nm*LTA and the SRL variant. (A) HPLC analysis of L-*threo*-4-$CH_3$-phenylserine and L-*erythro*-4-$CH_3$-phenylserine. The retention time of L-*threo*-4-$CH_3$-phenylserine and L-*erythro*-4-$CH_3$-phenylserine were 13.643 min and 20.932 min, respectively. (B) An ESI-MS spectrum of the product with $[M+H]^+$ at m/z 196.0973 and $[M+Na]^+$ at m/z 218.0790.

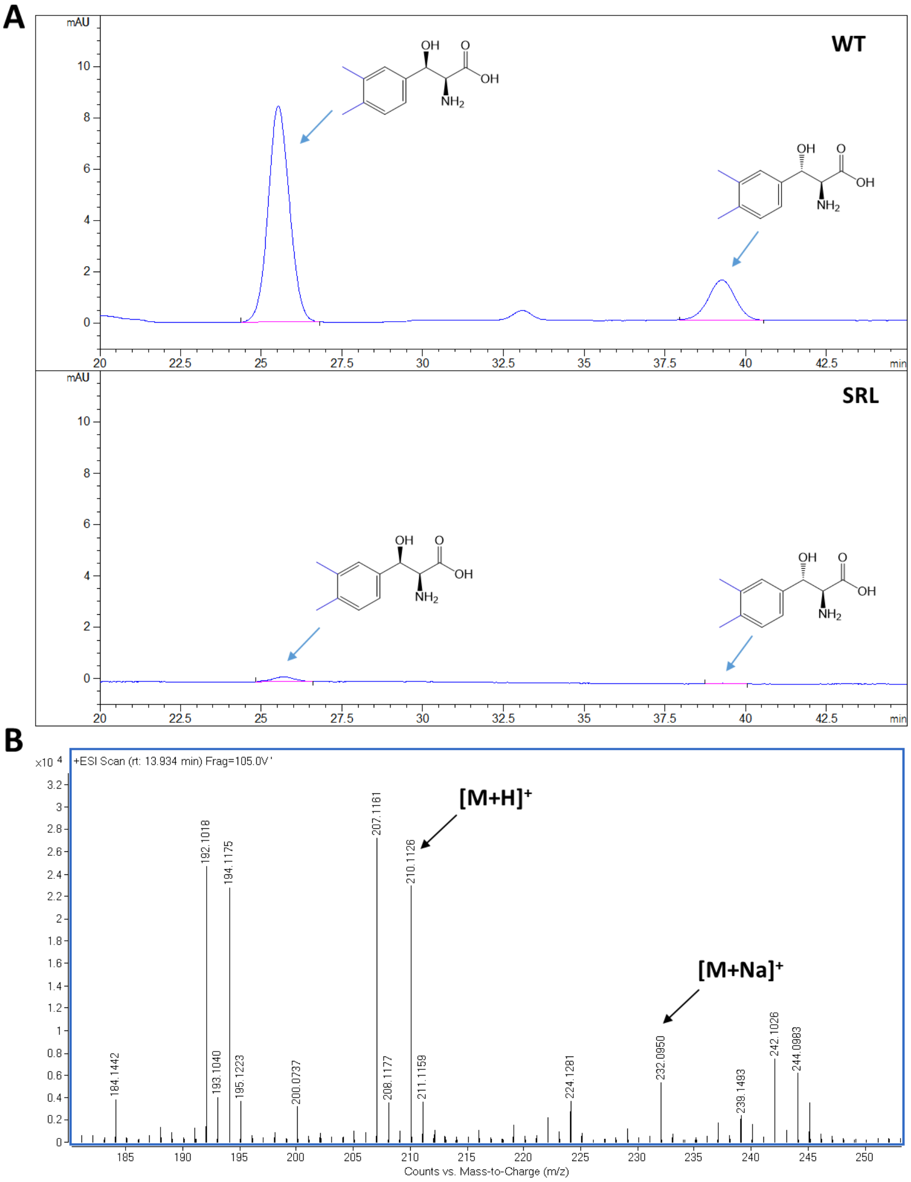

**Figure S27.** HPLC and MS analysis of L-3,4-$CH_3$-phenylserine catalyzed by WT *Nm*LTA and the SRL variant. (A) HPLC analysis of L-*threo*-3,4-$CH_3$-phenylserine and L-*erythro*-3,4-$CH_3$-phenylserine. The retention time of L-*threo*-3,4-$CH_3$-phenylserine and L-*erythro*-3,4-$CH_3$-phenylserine were 25.652 min and 39.294 min, respectively. (B) An ESI-MS spectrum of the product with $[M+H]^+$ at m/z 210.1126 and $[M+Na]^+$ at m/z 232.0950.

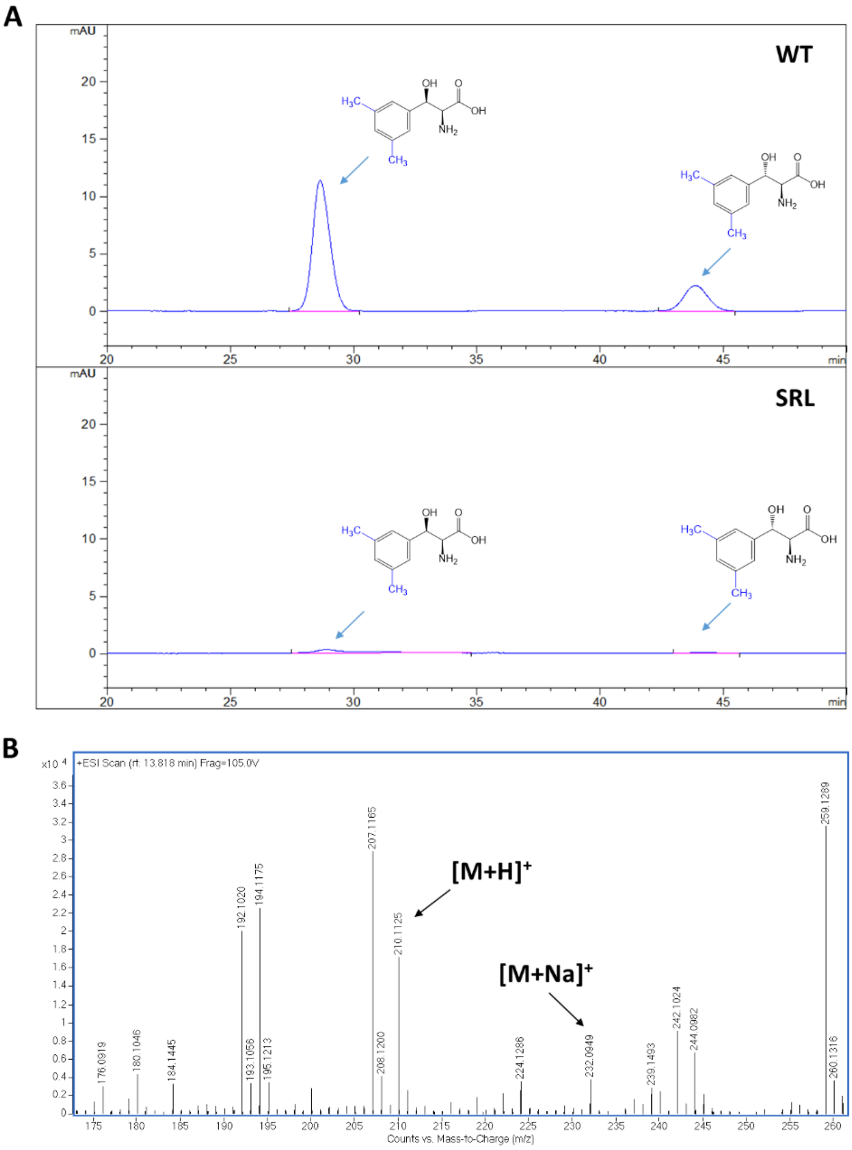

**Figure S28.** HPLC and MS analysis of L-3,5-$CH_3$-phenylserine catalyzed by WT *Nm*LTA and the SRL variant. (A) HPLC analysis of L-*threo*-3,5-$CH_3$-phenylserine and L-*erythro*-3,5-$CH_3$-phenylserine. The retention time of L-*threo*-3,5-$CH_3$-phenylserine and L-*erythro*-3,5-$CH_3$-phenylserine were 28.800 min and 43.892 min, respectively. (B) An ESI-MS spectrum of the product with $[M+H]^+$ at m/z 210.1125 and $[M+Na]^+$ at m/z 232.0949.

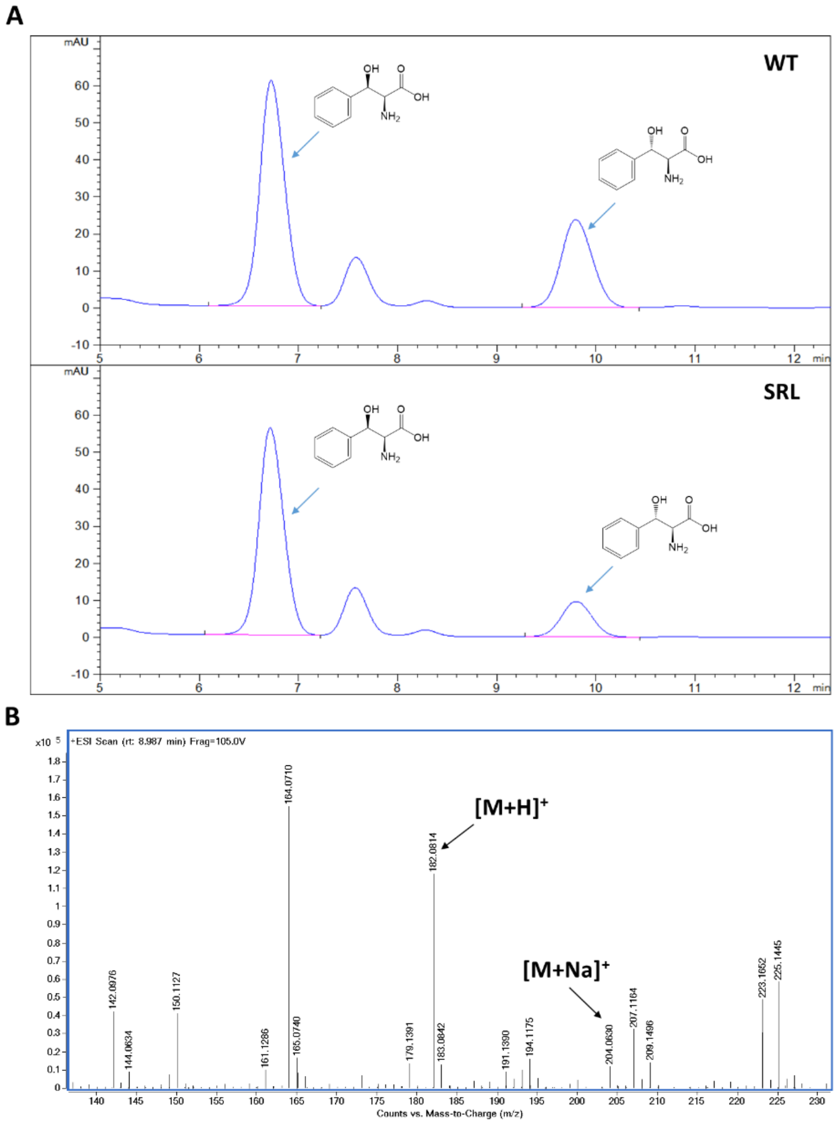

**Figure S29.** HPLC and MS analysis of L-phenylserine catalyzed by WT *Nm*LTA and the SRL variant. (A) HPLC analysis of L-*threo*-phenylserine and L-*erythro*-phenylserine. The retention time of L-*threo*-phenylserine and L-*erythro*-phenylserine were 6.722 min and 9.795 min, respectively. (B) An ESI-MS spectrum of the product with $[M+H]^+$ at m/z 182.0814 and $[M+Na]^+$ at m/z 204.0630.

### 1.6 Gene Sequences

Nucleotide and amino acid sequence for production of LTAs. All sequences are codon-optimized for expression in *E. coli*.

| ***Rp*LTA Nucleotide Sequence** |
| --- |
| ATGATATTTGCGTCAGATAATTGGGCAGGAGCTGCGGATGAGATCGCCGAAAGCCTGCGCCGCCA<br>CTCTGAAGGTTTTTCTCCGGCTTACGGTGAATCTCCGCTGGATAAGCAATTGGAAGCGAGATTCA<br>ACGATCTGTTCGAGCGTGAGGTGGCCGTTTTTTTGTTGGCACTGGTACCGCAGCGAATAGCCTC<br>GCTATGAGCGCATTTAACCGTCCGGGTGGTTTTGTTCTGTGTCATCGTGAGGCCCATATGATTGAG<br>GATGAGTGCGGTGCCCCCGAATTCTTCACCTCCGGCGCGCGTTTGGCTCCGATTGACGGCGCTTA<br>CGGTAAACTGGACCCAGAGCACCTGCGTGAGGGCCTTGAGCGCTTCGACCCGGGTTTTGTTCAT<br>CATGGTCAGCCTATGGCTGTGTCCTTGACGCAAGCGACCGAAGTCGGTACGGTGTATAGCTGCGA<br>TGAGCTGAAAGAAATCTCAGACCTGACCCACGCATTTGGCCTGCCGCTTCACATGGACGGCGCG<br>CGTTTCGCGAATGCAATGGTTCGTTTGGGAGTTAGCCCGGCGGAGATGACCTGGAAAGCGGGTG<br>TTGACATCCTGTCCTTCGGCGGTACGAAAAACGGCTGCTGGTGTGCAGAGGCGATTGTGTTCATG<br>GATCCGGCGAGGGCGAAGCAGCTGCCGTTTATCCGTAAACGCGCCGCACAGCTGTTTAGCAAAA<br>CCCGTTTCATCGCGGCACAATTTCACGCCTATCTGGACAACGACCTGTGGATTAGCCTGGCTAAG<br>CATTCCAACGCCATGTCGGATGAATTAGCGCGCCGCTTGGACCACTTCGAAGAACTGCGTATCGC<br>GTGGAAGTGCCAAAGCAATGAACTGTTCGTCACCATGCCGAAGGTGCTGGCTAAAAAGCTGCAC<br>GATCAGGGTGCAAAGTTCTACCCGTGGCCGGTGCCGGCGGAATTTGCGAGCAAGTTGCAGAAAG<br>GCGATGGCTTATATCGTTTCGTGACCAGCTTTGCGACCCAGCAAGAACAGATTGACGAACTTATC<br>GCGACCATTGAGGCAACTGTCGTGGCGTAA |
| ***Rp*LTA Amino Acid Sequence** |
| MIFASDNWAGAADEIAESLRRHSEGFSPAYGESPLDKQLEARFNDLFEREVAVFFVGTGTAANSLAMS<br>AFNRPGGFVLCHREAHMIEDECGAPEFFTSGARLAPIDGAYGKLDPEHLREGLERFDPGFVHHGQPM<br>AVSLTQATEVGTVYSCDELKEISDLTHAFGLPLHMDGARFANAMVRLGVSPAEMTWKAGVDILSFG<br>GTKNGCWCAEAIVFMDPARAKQLPFIRKRAAQLFSKTRFIAAQFHAYLDNDLWISLAKHSNAMSDE<br>LARRLDHFEELRIAWKCQSNELFVTMPKVLAKKLHDQGAKFYPWPVPAEFASKLQKGDGLYRFVTS<br>FATQQEQIDELIATIEATVVA |
| ***Nm*LTA Nucleotide Sequence** |
| ATGGCTAGTAATGATTCATGTATAGAAGACACAGTGAGCTTCACCTCCGACAACATCGCGGCTGC<br>GGCTCCGGAAATCGTGCAGGCGATGGCGCAAGCGTGTCAGGGCAATGCGCAACCGTACGGCGGA<br>GACGCGCTGACTCAAAATGTTGAGGCACAGCTTAAGGCTATCTTCGAGTGTGACCTCCAGCTGTT<br>CTTGGTACCGACGGGTTCGGCTGCCAACGCGATCAGCCTGGCTGCGCTCACCCCTCCGTGGGGG<br>GCGATTTTGTGCCACCAGGAGAGCCATATTAACAACGATGAGTGCGGTGCGCCGGAATTCTTTAC<br>CGCAGGTGCCAAACTGATCGCGGTGGCGGGCACCCATGGCAAACTGGATCCGCAGGCGTTGACT<br>CAAGCAGCGCGCAATAAACGCGGCGACGTTCACAGCGTCGAGCCGACCACCGTGAGTATTACCC<br>AGGCAACCGAAGTTGGTTCTATCTACGCGTTGGACGAGCTGAACGAGATTGGTCAGATTTGCCGT<br>AACGAAGGTCTGAAACTGCACATGGATGGTGCGCGTTTCGCTAACGCACTGTCTGCGCTGGGTT<br>GTACCCCAGCTGAAATGACCTGGAAGGCAGGCGTTGATGTGCTGAGCTTTGGTGCGACTAAGAA |

| |
|---|
| TGGTTCCCTTTGCGCCGAGGCTATCATCTTGTTCGATAAAAGCTATGCCCAAGAAATCGCGTTCCG<br>CCGTAAACGTGGTGGCCATCTGCTGTCCAAGATGCGTTTTCTCAGCGCACAAATGCATGCGTACC<br>TGGCGGACGACCTGTGGCTGACCAATGCCCGCCACGCGAATCTGATGGCAGCGCGTTTGGCTGC<br>TGGCCTGTCAGCCTTGAGCCGTGTTTCGCTGATCGCGCCGACCGAAAGCAACATTATCTTCTGCC<br>GCATGCCGACCAAGATGATTGCCGCATTACAGCAACAGGGTTTTCAGTTTTATCACGATCGTTGG<br>GGCGACGGCATTGTTAGACTGGTCACGTCTTTCGCCACGACGCAGGCTCAAGTGGATACCTTTAT<br>CGCGGCTGCCGCGCAACTGAACCAGAACACCGATTAA |
| ***Nm*LTA Amino Acid Sequence** |
| MASNDSCIEDTVSFTSDNIAAAAPEIVQAMAQACQGNAQPYGGDALTQNVEAQLKAIFECDLQLFLV<br>PTGSAANAISLAALTPPWGAILCHQESHINNDECGAPEFFTAGAKLIAVAGTHGKLDPQALTQAARNK<br>RGDVHSVEPTTVSITQATEVGSIYALDELNEIGQICRNEGLKLHMDGARFANALSALGCTPAEMTWK<br>AGVDVLSFGATKNGSLCAEAIILFDKSYAQEIAFRRKRGGHLLSKMRFLSAQMHAYLADDLWLTNA<br>RHANLMAARLAAGLSALSRVSLIAPTESNIIFCRMPTKMIAALQQQGFQFYHDRWGDGIVRLVTSFAT<br>TQAQVDTFIAAAAQLNQNTD |
| ***Sp*LTA Nucleotide Sequence** |
| ATGTTCTTTGCTTCAGATAATGCAGGACCCGTACACCCGCAGATTATGAATCGTCTGGCGCAAGCA<br>AACACCGGTCATGCAATGCCTTACGGCAATGATCCCATCATGGACGAAGTTCGTGATGCGATCCG<br>TACCGCCTTTGAAGCGCCGGAGGCCGCGGTGTACTTGGTTGCAACCGGCACGGCGGCGAACGCG<br>CTTGCGCTGGCGTGTTATACCCAGCCGTGGCAGACCATTTTCTGCAGCGTTACCAGCCACATCCA<br>CGAAGACGAGTGCAACGCTCCGGAGTTCTACGCAGGAGCGGCGAAATTGACCGTTGTCGAGAC<br>AGACGACAAAATGACGCCGGAGGCGCTGACGAAGGCCATCGAGAAACACCCGGAGGGTAACGT<br>ACACGGCGCCCAACGTGGTCCGGTTAGCATCACCCAAGTTACCGAAAGAGGCTCCGTTCACACC<br>CTCGAGGAATTGAATGCACTCACCGCTGTGGCGAAAAGCTATGATCTGCCGGTGCACCTTGACGG<br>CGCTCGCTTCGCAAACGCGTTGGTCGCGCTGGGTTGCACCCCGGCGGAAATGACTTGGAAGGCA<br>GGGGTTGATGTGGTGTCTTTTGGTGGTACGAAAAACGGCTGTATGGGCGTGGAAGCGGTCATCTT<br>CTTTGATCCGGCGAAGGCATGGGAATTCGAGCTGCGTCGCAAGCGTGGTGCGCATCTGTTCAGCA<br>AGCATCGCTTTCTGTCCGCCCAAATGGCTGGTTATATGCAGGATGACCTGTGGAAAACCACCGCT<br>GCGCGCGCGAACGCGAACGCGCGTCATTTAGCTGAGGGTCTGCGCACCGCTGGTGCGACGCTGC<br>TGCACAAACCAGATGCAAACATGATTTTCGCCAAGTGGCCGCGTCGTATTCACCAGAAGCTGCAT<br>GATGCAGGCGCCAAGTACTACGTGATGGACGGTCCGTTGGAAGGCGACGACCCAAATGAACCGC<br>TGTCAGCCCGTCTGGTGTGCGACTGGTCTATCGGCACTGAGGCGATTGATCAGTTTTTGTCGCATT<br>TTTAA |
| ***Sp*LTA Amino Acid Sequence** |
| MFFASDNAGPVHPQIMNRLAQANTGHAMPYGNDPIMDEVRDAIRTAFEAPEAAVYLVATGTAANAL<br>ALACYTQPWQTIFCSVTSHIHEDECNAPEFYAGAAKLTVVETDDKMTPEALTKAIEKHPEGNVHGAQ<br>RGPVSITQVTERGSVHTLEELNALTAVAKSYDLPVHLDGARFANALVALGCTPAEMTWKAGVDVVSF<br>GGTKNGCMGVEAVIFFDPAKAWEFELRRKRGAHLFSKHRFLSAQMAGYMQDDLWKTTAARANAN<br>ARHLAEGLRTAGATLLHKPDANMIFAKWPRRIHQKLHDAGAKYYVMDGPLEGDDPNEPLSARLVC<br>DWSIGTEAIDQFLSHF |
| ***Ml*LTA Nucleotide Sequence** |

| |
|---|
| ATGAACTTTGCTTCAGATAATGCAAGTCCCGTACCGCAGCAGGTTCTGGACGTGCTCGCCCGCGT<br>GAACTCCGGTGCGGCAGCCTCCTATGGTGCGGACGACGTAACGGCAGAGGTTGCGGATCGTGTT<br>CGTGCCTTGTTTGAAGCGCCAGGTGCGGCGGTGTACCTGGTGGCAACCGGTACAGCTGCGAACA<br>GCCTGTCCCTGGCGACTTTGTGCGCACCGTTTCAGACCATTTTCTGCAGCGAACATGCACACATC<br>CACGAAGACGAGTGCAATGCGCCGGAGTTCTATACCGGAGGTGCTAAGCTGACCTTGGTGCGTG<br>GCGGCGATGTTATGACTCCGGAGGCGTTGCGTAGCGCAATTCTGGGTGAAGGCAACCGCGGCGT<br>GCACGGCCCGCAGCGTGGTCCGGTTAGCGTTACGAATGTTACGGAGGGTGGCAATGTGTACGCG<br>TTGTCTGATATCGGTGCCTTATGCGCCGTTGCCCGCGAATATGGCTTACCGGTGCACTTGGACGGC<br>GCTCGCTTTGCAAACGCGTGTGTCAAGCTGGGGTGCACCCCAGCTGAGATGACCTGGAAAGCTG<br>GCGTCGATATTGCAGTGTTCGGCGGTACGAAAAACGGTCTGATGGACGCAGAGGCGGTCGTGAT<br>CTTCGATCCGGAGGCTCCGGCGAGCTCTGGTTTTACCCGTGCGCAAGAATTCGAGCTGCGCGTCA<br>AACGTGCAGGCCATCTGTACAGCAAGCATCGTTATGTTGCGGCACAGATGCTGGCGTACCTGGAA<br>GACGACCTGTGGAGAGATCTGGCGCAACAAGCGAATGATCATTGTGAAACCTTAGCTCGTGGCC<br>TGCAGGATATGGGTTTGGAGATCGTTAATAAGACCCGTGCGAACATGCTGTTCTTTCGTGCTCCGC<br>TGCGCGCGCACAAGGCGGCTCAAGCAGCGGGTGCCGTGTACGCGTTGTGGGGTAACCCGCCTGA<br>AAAAGCCGACGAACCGGCGCTGGCGCGCCTCGTTTGTAACTGGTCGACCACCGAGGCAGAAATC<br>ACCGAGTTCCTGAAAGTTATGCGTGCTGCATTGTAA |
| ***Ml*LTA Amino Acid Sequence** |
| MNFASDNASPVPQQVLDVLARVNSGAAASYGADDVTAEVADRVRALFEAPGAAVYLVATGTAANSL<br>SLATLCAPFQTIFCSEHAHIHEDECNAPEFYTGGAKLTLVRGGDVMTPEALRSAILGEGNRGVHGPQR<br>GPVSVTNVTEGGNVYALSDIGALCAVAREYGLPVHLDGARFANACVKLGCTPAEMTWKAGVDIAVF<br>GGTKNGLMDAEAVVIFDPEAPASSGFTRAQEFELRVKRAGHLYSKHRYVAAQMLAYLEDDLWRDLA<br>QQANDHCETLARGLQDMGLEIVNKTRANMLFFRAPLRAHKAAQAAGAVYALWGNPPEKADEPALA<br>RLVCNWSTTEAEITEFLKVMRAAL |
| ***Al*LTA Nucleotide Sequence** |
| ATGCATATGTATTTTGCTTCTGATAATTCAAGTCCCGTACCTCCGCAGATTTTGGACGCGTTGGTTC<br>ACGCAAACCATGGTTACGCTATGCCGTATGGTGCCGACACGATCATGGACAGCGTGCGTAACAAA<br>ATCCGCGAAGTGTTCGAGGCGCCAGAAGCGGCGGTGTATTTGGTTCCGACCGGCACCGCAGCGA<br>ACGTCTTGGCACTGAGCTGCCTGTGCCCGCCGTGGGCGACGATCTACTGCCATCAGAATGCTCAC<br>ATCGAGGAGGACGAGTGCGGTGCGCCGGAGTTCTATACCGGTGGCGCGAAGTTGACCCTGGTCG<br>GCGGTGACGACGCGAAAATGAGCCCGGAAGCGCTGAAGCAGGCAATTAGCTTTACTGCACGCGC<br>AGGCGTGCACAATGTGCAGAAAGGCGCGGTTAGCATTACCAATATCACCGAAAACGGCGCCCTG<br>TACTCTGCTGATGAGGTTCGTGCCTTATGCGATATTGCAAAGGCAAGCGATTTGCCGGTCCACATG<br>GACGGTGCGCGTTTTGCCAACGCCGTTGTTGGTGCCGGTTGTACCCCGGCAGAAATGACCTGGA<br>AGGCGGGCGTGGATGTACTTTCCTTTGGTGGTACGAAAAACGGCCTGATGGGTGTCGAGGCTGT<br>GGTTCTGTTCGATCCGAAGCGTGCTTGGGAGTTCGAGTTACGTCGTAAACGTGGAGGCCATCTGT<br>TTTCTAAGCACCGCTACCTGTCCGCGCAAATGGATGCCTATCTGGAGGATGACCTGTGGCTGAAA<br>CTGGCGACTCGCGCTAACGACGCGGCGGCGAGACTATCGAAAGGTATTTTGACCATCGAAGGCG<br>CTTCGCTGCTGCATCCGACCGACGGTAATGCGGTTTTCGCACGCTGGCCACGTGAAGGCCACCGT |

| |
|---|
| AGAGCGCAAGATGCGGGCGCTGTGTACTACCTGTGGCCGATGAACCAAAGCCTGGAGGGTCCGG<br>ATGAAGAACCGCTGTCCGCGCGTCTCGTGTGTTCTTGGTGTACCAGCAGCGCTGACGTTGAAAA<br>GTTTCTGGAACTGATCCGTGGTCTCGAGTAA |
| ***Al*LTA Amino Acid Sequence** |
| MYFASDNSSPVPPQILDALVHANHGYAMPYGADTIMDSVRNKIREVFEAPEAAVYLVPTGTAANVLA<br>LSCLCPPWATIYCHQNAHIEEDECGAPEFYTGGAKLTLVGGDDAKMSPEALKQAISFTARAGVHNVQ<br>KGAVSITNITENGALYSADEVRALCDIAKASDLPVHMDGARFANAVVGAGCTPAEMTWKAGVDVLS<br>FGGTKNGLMGVEAVVLFDPKRAWEFELRRKRGGHLFSKHRYLSAQMDAYLEDDLWLKLATRAND<br>AAARLSKGILTIEGASLLHPTDGNAVFARWPREGHRRAQDAGAVYYLWPMNQSLEGPDEEPLSARLV<br>CSWCTSSADVEKFLELIRG |
| ***Pp*LTA Nucleotide Sequence** |
| ATGACAGATAAAAGCCAGCAGTTTGCAAGCGATAATTATAGCGGCATTTGCCCAGAAGCATGGGC<br>AGCAATGGAAAAGGCAAACCACGGTCATGATCGCGCATACGGTGATGATCAGTGGACCGAACGT<br>GCAAGCGAATATTTTCGTAATCTGTTTGAAACAGATTGCGAAGTTTTCTTCGCATTTAACGGTACA<br>GCAGCCAACAGCCTGGCACTGGCAAGCCTGTGTCAGAGCTATCATAGCGTTATTTGTAGCGAAAC<br>CGCACATGTTGAAACAGATGAATGTGGTGCACCTGAATTTTTCAGCAATGGCAGTAAACTGCTGA<br>CCGCAGCAAGTGTGAATGGCAAACTGACCCCGCAGAGTATCCGTGAAGTGGCACTGAAACGTCA<br>GGATATTCATTATCCGAAACCTCGTGTTGTTACCATCACACAGGCAACCGAAGTTGGTACAGTTTA<br>TCGCCCGGATGAACTGAAAGCAATTAGCGCAACATGTAAAGAACTGGGTCTGAATCTGCACATG<br>GATGGCGCACGTTTTACCAATGCGTGCGCATTTCTGGGTTGTAGTCCAGCAGAACTGACCTGGAA<br>AGCTGGCGTTGATGTTCTGTGTTTTGGTGGTACTAAAAATGGCATGGCAGTTGGTGAAGCAATTC<br>TGTTCTTCAATCGTCAGCTGGCAGAAGATTTTGATTATCGTTGTAAACAGGCAGGTCAGCTGGCA<br>AGTAAAATGCGTTTTCTGAGCGCACCGTGGGTTGGTCTGCTGGAAGATGGTGCCTGGCTGCGTCA<br>TGGTAATCATGCAAATCATTGTGCCCAGCTGCTGGCACTGCTGGTTAGTGATCTGCCTGGTGTTGA<br>ACTGATGTTTCCGGTTGAAGCAAATGGTGTTTTCCTGCAGATGCCGGAACATGCAATCGAAGCGC<br>TGCGTGCGAAAGGTTGGCGTTTTTATACTTTTATTGGTAGCGGTGGTGCACGTTTTATGTGTAGCT<br>GGGATACCGAAGAAGAACGTGTTCGTGAACTGGCAGCAGATATTCGTAGCATTATTACAGCATAA |
| ***Al*LTA Amino Acid Sequence** |
| MTDKSQQFASDNYSGICPEAWAAMEKANHGHDRAYGDDQWTERASEYFRNLFETDCEVFFAFNGT<br>AANSLALASLCQSYHSVICSETAHVETDECGAPEFFSNGSKLLTAASVNGKLTPQSIREVALKRQDIHY<br>PKPRVVTITQATEVGTVYRPDELKAISATCKELGLNLHMDGARFTNACAFLGCSPAELTWKAGVDVL<br>CFGGTKNGMAVGEAILFFNRQLAEDFDYRCKQAGQLASKMRFLSAPWVGLLEDGAWLRHGNHAN<br>HCAQLLALLVSDLPGVELMFPVEANGVFLQMPEHAIEALRAKGWRFYTFIGSGGARFMCSWDTEEE<br>RVRELAADIRSIITA |
| ***Aj*LTA Nucleotide Sequence** |

ATGCGTTATATCGATCTGCGTAGCGATACCGTTACCCAGCCTACCGATGCAATGCGTCAGTGTATGC
TGCATGCAGAAGTTGGTGATGATGTTTATGGTGAAGATCCTGGTGTTAATGCACTGGAAGCCTATG
GTGCAGATCTGCTGGGTAAAGAAGCAGCACTGTTTGTTCCGTCTGGTACGATGAGCAATCTGCTG
GCAGTTATGAGCCATTGTCAGCGTGGTGAAGGTGCCGTTCTGGGTAGTGCAGCACATATTTATCGT
TATGAAGCACAGGGTAGTGCCGTTCTGGGCAGTGTTGCACTGCAGCCTGTTCCGATGCAGGCAG
ATGGTAGCCTGGCGCTGGCTGATGTTCGTGCTGCAATTGCACCGGATGATGTTCATTTTACCCCAA
CCCGCCTGGTTTGTCTGGAAAATACCCATAATGGTAAAGTCCTGCCACTGCCGTATCTGCGTGAA
ATGCGTGAACTGGTTGATGAACATGGTCTGCAGCTGCATCTGGATGGCGCACGTCTGTTTAATGC
CGTTGTGGCAAGCGGTCATACAGTGCGTGAACTGGTGGCACCGTTTGATAGCGTTTCTATTTGTCT
GAGCAAAGGTCTGGGTGCACCGGTTGGTAGCCTGCTGGTTGGTAGCCATGCATTTATTGCACGTG
CACGTCGCCTGCGTAAAATGGTTGGTGGTGGTATGCGTCAGGCCGGTATTCTGGCCCAGGCTGGT
CTGTTTGCCCTGCAGCAGCATGTTGTTCGTCTGGCCGATGATCATCGTCGTGCACGTCAGCTGGC
AGAAGGTCTGGCAGCTCTGCCTGGTATTCGTCTGGATCTGGCACAGGTTCAGACCAATATGGTTT
TCCTGCAGCTGACCTCTGGTGAAAGCGCACCGCTGCTGGCATTTATGAAAGCACGTGGTATTCTG
TTTTCTGGTTATGGTGAACTGCGTCTGGTTACCCATCTGCAGATTCATGATGATGATATTGAAGAA
GTGATCGATGCATTTACCGAATATCTGGGTGCATAA

***Aj*LTA Amino Acid Sequence**

MRYIDLRSDTVTQPTDAMRQCMLHAEVGDDVYGEDPGVNALEAYGADLLGKEAALFVPSGTMSN
LLAVMSHCQRGEGAVLGSAAHIYRYEAQGSAVLGSVALQPVPMQADGSLALADVRAAIAPDDVHFT
PTRLVCLENTHNGKVLPLPYLREMRELVDEHGLQLHLDGARLFNAVVASGHTVRELVAPFDSVSICLS
KGLGAPVGSLLVGSHAFIARARRLRKMVGGGMRQAGILAQAGLFALQQHVVRLADDHRRARQLAE
GLAALPGIRLDLAQVQTNMVFLQLTSGESAPLLAFMKARGILFSGYGELRLVTHLQIHDDDIEEVIDA
FTEYLGA

***Ec*LTA Nucleotide Sequence**

ATGATCGATCTGCGCAGCGATACCGTTACCCGCCCTAGTCGTGCAATGCTGGAAGCAATGATGGC
AGCCCCGGTTGGTGATGATGTTTATGGCGATGATCCGACCGTTAACGCACTGCAGGATTATGCTGC
AGAACTGAGCGGTAAAGAAGCAGCAATTTTCCTGCCGACCGGTACACAGGCAAATCTGGTAGCA
CTGCTGAGTCATTGTGAACGTGGTGAAGAATATATTGTTGGTCAGGCCGCACATAATTATCTGTTT
GAAGCAGGTGGTGCAGCCGTTCTGGGTAGTATTCAGCCGCAGCCTATTGATGCCGCAGCAGATGG
TACACTGCCGCTGGATAAAGTTGCAATGAAAATTAAACCGGATGATATTCATTTTGCCCGTACAAA
ACTGCTGAGCCTGGAAAACACCCATAACGGCAAAGTTCTGCCGCGTGAATATCTGAAAGAAGCA
TGGGAATTTACCCGCAAACGCAACCTGGCACTGCATGTTGATGGTGCACGCATCTTTAATGCAGT
TGTAGCATACGGTTGTGAACTGAAAGAAATCACCCAGTATTGTGATAGCTTTACCATTTGTCTGAG
CAAAGGTCTGGGTACACCGGTTGGCAGCCTGCTGGTTGGTAATCGTGATTATATTAAACGCGCCAT
TCGTTGGCGCAAAATGACCGGTGGTGGTATGCGCCAGAGCGGTATTCTGGCAGCAGCAGGTATGT
ATGCACTGAAAAACAATGTTGCACGTCTGCAGGAAGATCATGATAATGCAACCTGGATGGCAGAA
CAGCTGCGTGAAGCAGGTGCAGATGTGATGCGTCAGGATACCAATATGCTGTTTGTTCGTGTGGG
CGAAGAAAATGCAGCAGCACTGGGTGAATATATGAAAGCACGTAACGTACTGATTAATGCAAGCC
CGATTGTTCGTCTGGTTACCCATCTGGATGTTAGCCGTGCCCAGCTGGCAGAAGTGGCAGCACAT
TGGCGTGCTTTTCTGGCCCGTTAA

| ***Ec*LTA Amino Acid Sequence** |
| --- |
| MIDLRSDTVTRPSRAMLEAMMAAPVGDDVYGDDPTVNALQDYAAELSGKEAAIFLPTGTQANLVA LLSHCERGEEYIVGQAAHNYLFEAGGAAVLGSIQPQPIDAAADGTLPLDKVAMKIKPDDIHFARTKLL SLENTHNGKVLPREYLKEAWEFTRKRNLALHVDGARIFNAVVAYGCELKEITQYCDSFTICLSKGLG TPVGSLLVGNRDYIKRAIRWRKMTGGGMRQSGILAAAGMYALKNNVARLQEDHDNATWMAEQLR EAGADVMRQDTNMLFVRVGEENAAALGEYMKARNVLINASPIVRLVTHLDVSRAQLAEVAAHWR AFLAR |
